# Successful Predictive Modeling of Pollen Fitness Phenotypes Is Enabled by Measures of Expression Specificity

**DOI:** 10.64898/2025.12.05.692487

**Authors:** Sebastian A.F. Mueller, Zuzana Vejlupkova, Molly Megraw, John E. Fowler

## Abstract

The ability to predict phenotypes from genotypes in multicellular organisms remains limited despite rapid advances in genotyping and phenotyping methods. Machine learning offers a promising way to model phenotype from genotype, but requires sizable datasets that quantitatively link phenotype to specific genes. Such datasets remain limited; however, maize pollen provides a unique biological system that is especially well suited for this challenge. Because maize pollen is haploid, mutations that affect its function can result in a quantitative phenotypic effect on pollen fitness, measurable as deviations in transmission rate from the expected Mendelian ratio. We leveraged a large set of fluorescently-marked insertional mutations, the *Ds-GFP* lines, to link fitness effects to specific genes. We then developed a machine learning framework that integrates expression profiling and genomic data to predict genes contributing to pollen fitness in maize. Well performing models that distinguish genes with strong fitness effects from those with little or no fitness effect could be generated using features, such as codon usage, derived solely from genome sequence (auROC 85%). Using expression data enabled even more successful models, achieving auROC values above 90%. Because we used interpretable machine learning methods, we were able to identify expression specificity as a critical feature for strongest model performance. The best performing model was achieved when specificity measures were complemented with certain genomic sequence features. Models that include expression specificity generalize well across the maize genome, as predictions meet expectations of mutational frequencies for thousands of genes in a well characterized mutagenized population.

## 1 Introduction

Despite the rapid advances in genotyping (Farooq et al., 2024; Bansal and Srivastava, 2018; Younessi-Hamzekhanlu et al., 2022) and phenotyping (Visakh et al.; Wan et al., 2025) methods, our ability to predict phenotypes from genotypes in multicellular organisms remains limited (Baxter, 2020). Because phenotypes arise in part from complex interactions among genotypically determined biological systems, models must capture this complexity by extracting patterns that link genomic factors to quantifiable phenotypes (Amin et al., 2025). Machine learning (ML) is well suited to this challenge, as it can integrate disparate datatypes and generate interpretable insights.

ML models have contributed to pioneering studies that investigated the use of systems-level gene properties to predict phenotypic outcomes. For example, an ML model built from 23 properties was able to predict haploinsufficient phenotypes for genes in *S. cerevisiae*, with higher likelihood correlating with measures of protein-protein interaction, sequence conservation, and proteomic abundance (Norris et al., 2013). Subsequent human studies demonstrated that applying richer model architectures to epigenomic data improved both performance and generalizability in attempts to predict genes that show phenotypes with only one mutant copy (Yang et al., 2021). More recently, models such as QTG-Finder have been used to help prioritize potential for causality out of a limited number of genes that lie within identified Quantitative Trait Loci intervals in rice (Lin et al., 2019, 2020; Hartanto et al., 2022). Genome wide-association studies (GWAS) provide another quantitative approach to explore genome to phenome relationships and have been used extensively in multicellular eukaryotes (Huang and Han, 2014; Xiao et al., 2017; Tibbs Cortes et al., 2021; Uffelmann et al., 2021). While these studies advance our understanding of genotype to phenotype relationships, the GWAS approach typically involves natural populations, which are limited in the types and extent of genetic variation that can contribute to phenotypes for study. Thus, GWAS can only implicate specific loci, variants, or gene sets as linked to particular phenotypes.

Studies employing ML modeling have shown that certain omics-scale data can be used successfully to predict lethal phenotypes (Lloyd et al., 2015) and complex plant traits in Arabidopsis accessions (e.g., flowering time) (Wang et al., 2024), or to associate Gene Ontology (GO) terms with gene models in maize (Dai et al., 2020). Yet, there have been calls for models more specifically capable of predicting whether a gene influences a given phenotype, particularly in plants (Schnable, 2020). Despite the helpful advancements discussed above, this goal remains elusive (Baxter, 2020). A model that predicts phenotypic outcomes on a gene by gene basis would help advance a global mechanistic understanding of multicellular biological systems. However, even with the abundance and availability of sophisticated omics-scale data, this task remains challenging. Difficulties arise because many gene models cannot be easily connected with phenotypes, phenotypic measures are difficult to standardize across all possible outcomes, and gathering data to confidently conclude that alteration of a single gene function has no effect on phenotypes is often impractical. Because gene models associated with distinct phenotypes may differ substantially in their underlying features, specific genotype-phenotype relationships may be obscured in more generalized datasets (Schnable, 2020). Thus, modeling single gene contributions requires datasets in which specific phenotypic outcomes are quantified for many genes. Such datasets enable models to learn complex relationships between gene level omics data and phenotypic outcomes.

The maize male gametophyte is a unique system that is especially well suited to addressing these challenges, with several characteristics that enable efficient quantitative phenotyping. Although it is a multicellular system, it is comparatively simple, consisting of three cells at maturity. Two of these are sperm cells, which are essential for the gametophyte’s function: to produce the next generation (the seed, i.e. the kernel in maize) via double fertilization. Because it carries only a single copy of its genome (haploid), mutations are expressed in the first generation, eliminating the need for further genetic crosses to reveal any phenotypic effects. Furthermore, the structure of the maize ear, which encompasses all kernels from a controlled pollination, enables statistically robust high-throughput phenotyping. That is, the outcomes of many double fertilizations are gathered into single ears, where each kernel reflects the reproductive success of a genetically independent male gametophyte. Because an ear bears hundreds of kernels in an orderly arrangement, kernel counts within a ear provide a useful measure of the fitness effect for each mutation. In practice, these biological characteristics can be exploited for large scale dataset generation of quantitative phenotypes with the *Ds-GFP* mutant population (Garcia et al., 2017; Li et al., 2013; Xiong et al., 2013). This resource includes thousands of lines carrying single gene insertional mutations tagged with a marker visualized as a fluorescent kernel. Together, these elements make systematic *Ds-GFP* insertional mutagenesis a powerful approach for quantitatively linking individual genes to their phenotypic outcomes.

A pilot study (Warman et al., 2020) used this approach to assess the phenotype of mutations in 52 pollen genes, finding a correlation between high levels of transcripts of the wild type gene and loss of pollen fitness when the gene is mutated. Here we used a much expanded dataset with 270 genes to develop machine learning models that can successfully predict whether loss of a particular gene’s function strongly reduces pollen fitness. Our modeling process explored hundreds of genomic, transcriptomic, and proteomic features to predict loss-of-function phenotypes in maize pollen genes. Using interpretable modeling methods, we were able to identify the datatypes that are most predictive of phenotype. Notably, this analysis uncovered distinctive expression characteristics that enabled strong predictive model performance that is likely to generalize well to the whole genome. Furthermore, genomic features can be integrated into the model to further increase predictive power beyond what could be achieved with expression information alone.

## 2 Results

### 2.1 Unique Pollen Dataset Enables Fitness Modeling

Guided by publicly available expression data (Warman et al., 2020; Walley et al., 2016; Woodhouse et al., 2021b), we used the EarVision phenotyping system (Ruggiero et al., 2025) to generate the largest possible training dataset linking pollen-expressed gene models to a fitness measure associated with insertional mutations in those gene models. Because these fitness measurements represent the final outcome of competition between mutant and wild-type pollen (i.e., success in generating progeny kernels), they should capture a broad range of cellular-level defects, providing a comprehensive phenotypic data set. First, all gene models associated with above-background expression levels in pollen (including sperm cells), either by RNA-seq or proteomic profiling, were identified. By cross-referencing this set of gene models with the genomic locations of *Ds-GFP* insertions (Xiong et al., 2013), more than 400 insertion alleles predicted to contain a disruption in the coding sequence of pollen-expressed gene models were obtained. The locations of 305 of these insertions were validated using allele-specific primers and Sanger sequencing (Methods, *SI Appendix*, Data S1), and then targeted for pollen fitness measurement. Plants heterozygous for wild-type and mutant *Ds-GFP* alleles for each targeted insertion were then used as pollen parents in an outcross to a wild-type ear, with the expectation of 1:1 Mendelian segregation of the resulting kernels if the mutation had no effect on pollen fitness. For alleles that are deleterious to pollen fitness, direct competition between mutant and wild-type gametophytes upon pollination results in a transmission rate of less than 50%, which can be quantitatively measured as the ratio of GFP to non-GFP seeds on each ear (Fig. 1A). Over 3000 such ears were assessed using EarVision, providing a standardized method to phenotype and count over one million kernels (*SI Appendix* Data S2) (Fig. 1B), which we then used to estimate relative transmission rates for each mutant allele with data from at least two independent ears (*SI Appendix*, Data S3) (Fig. 1C).

**Figure 1.**
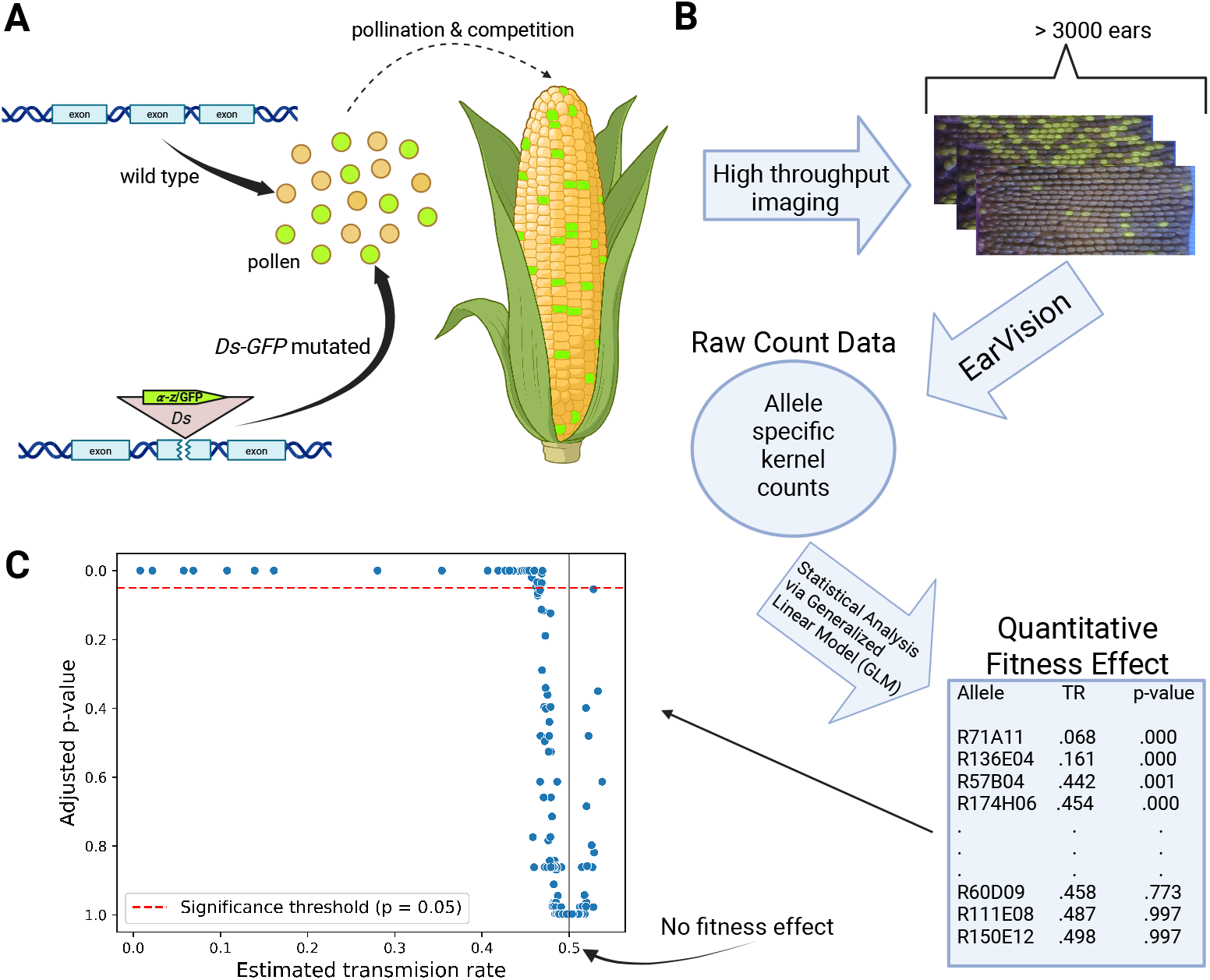
Quantifying pollen fitness effects of *Ds-GFP* insertions using the EarVision system (Ruggiero et al., 2025). *(A)* Heterozygous plants carrying *Ds-GFP* insertions were outcrossed to wild-type, placing mutant and wild-type pollen in direct competition during pollination. Assuming equal fitness, the expected ratio of progeny kernels is 1:1, yielding approximately half wild-type and half fluorescent (GFP marked) progeny kernels. Reduced frequency of GFP kernels indicates that the insertion is associated with reduced pollen fitness. *(B)* The EarVision system (Ruggiero et al., 2025) was used to image over 3,000 different ears. Kernel counts were analyzed using a generalized linear model (GLM) to estimate allele specific transmission rates. Only alleles with PCR validated *Ds-GFP* insertion sites and data from two or more ears were used. *(C)* Adjusted p-values are plotted against the estimated transmission rates for all 305 validated alleles. The majority of alleles are associated with a transmission rate of 0.5, indicating No fitness effect. Figure created in BioRender. Mueller, S. (2025) https://BioRender.com/ld96nl1

Initial assessment of these data revealed that three-quarters (28/37) of the alleles with significant defects (adj p <0.05) were associated with transmission rates in the 40%-47% range. This trend is consistent with expectations, as the *Ds-GFP* population was generated with mutagenized genomes transmitted through pollen (Li et al., 2013), reducing the likelihood of recovery of alleles with severe effects. Alleles with transmission rates below 45% were classified as “strong fitness effect” as all had small adjusted p-values (p <.001), indicating a significant deviation from the expected 1:1 ratio. Conversely, alleles with transmission rates above 48% all consistently had non-significant adjusted p-values, supporting their classification as “no fitness effect”. Alleles with transmission rates between 45%-48% were excluded from modeling because this range represents a biologically ambiguous zone in which neither statistical tests nor biological interpretation provide a clear distinction between mildly defective and no effect outcomes. While the chosen boundaries of 45%-48% are somewhat heuristic, this conservative design choice minimizes potential labeling noise and ensures a more robust classification scheme for downstream modeling. After mapping the validated insertion sites to B73v5 genome annotation, 15 alleles not associated with canonical transcript models were excluded. Insertion sites for 97% of the remaining alleles interrupted coding sequence (CDS) or the ‘5 untranslated region (5’UTR) of the canonical transcript models, strongly suggesting that the phenotypic classifications represent the consequence of reduced function for each associated gene model. Further evidence for this conclusion is that congruent phenotypes were present for independent alleles interrupting the nine non-excluded gene models represented by multiple insertions (two “strong fitness effect”, seven “no fitness effect”, *SI Appendix*, Data S4). For such alleles, the one with the most severe effect was used as representative, to assign a fitness phenotype to the gene model. With these exclusions, our final high-confidence dataset for modeling pollen fitness effects encompassed 227 unique gene models with pollen fitness measurements. Warman et al. (2020) previously noted that gene expression level correlates with the severity of transmission defect after mutation. In the following section, we explore this correlation more deeply using a greatly expanded set of omics-scale data types.

### 2.2 Multiple Genomic Feature Sets Can Generate Successful Pollen Fitness Models

For each of the 227 unique gene models with high-confidence pollen fitness measurements gathered above, we collected publicly available expression profiling and genomic feature data (Sen et al., 2023; Walley et al., 2016; Hufford et al., 2021). Specifically, over 600 distinct data features associated with each gene model were collected, including mRNA levels, protein levels, DNA base composition, amino acid composition properties, synteny with orthologs, KaKs ratio, and epigenomic features such as DNA methylation. We combined these data to create a ground truth data set of outcomes and associated features for all of our data points (i.e., gene models) (*SI Appendix*, Data S5).

To assess the feasibility of accurate modeling, we trained and tested machine learning models using multiple distinct sets of input features. For each feature set, we trained a logistic regression model and evaluated performance using the average auROC across all test folds from 3-fold cross-validation. In the general modeling framework (*SI Appendix*, Fig. S1), different features are used as model inputs to predict a phenotype of either strong fitness effect or no fitness effect for a gene model. Using an iterative modeling process, we developed three successful model types: (1) a Genomic Model, using only features derived from genome sequence; (2) an Expression Model, using only RNA and proteomic profiling features; and (3) a Combination Model using both expression and genomic sequence features. The complete list of features used for each model can be found in *SI Appendix*, Table S1. Model performance metrics are summarized in Figure 2. These results demonstrate that successful predictive models of pollen fitness outcome (auROC > 85%) can be constructed using our high-confidence quantitative phenotypic dataset. As a result of this predictive success, the modeling process can be interpreted to help identify the features that are most informative for pollen fitness outcomes.

**Figure 2.**
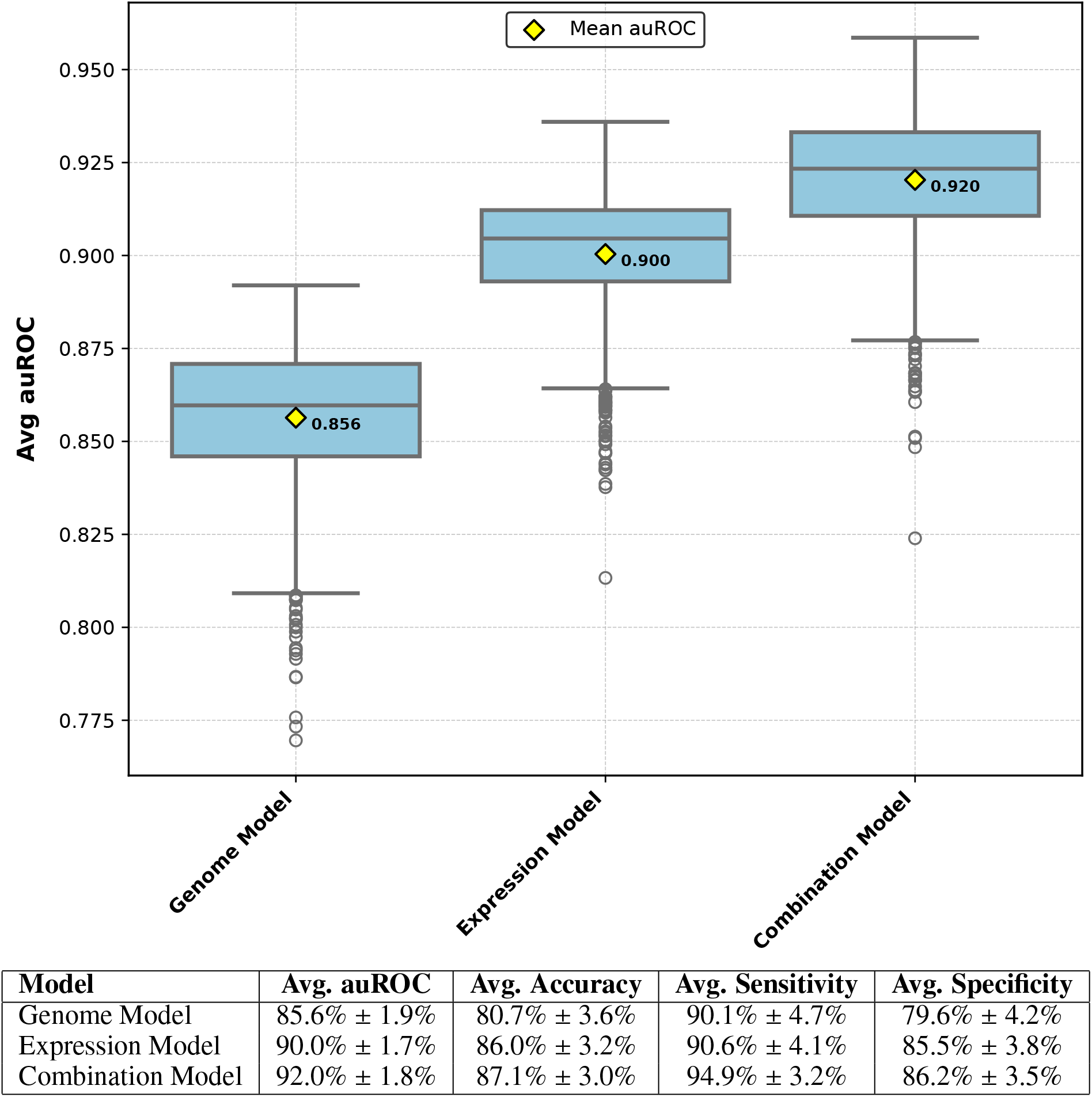
Strong predictive performance for fitness phenotype classification can be achieved using diverse feature sets. The Genome Model uses only genomic sequence derived features, the Expression Model uses only RNA-Seq and proteomic profiling features, and the Combination Model integrates both feature types (see *SI Appendix*, Table S1). All reported metrics represent the average test fold auROC across 1,000 cross-validation seeds, reported with their standard deviations. Among all the models, the Combination Model achieved the highest performance across every metric, including auROC, accuracy, sensitivity and specificity.

### 2.3 RNA and Protein Expression Data Sets Produce Similar Model Performance

Our final models used two types of expression data: RNA sequencing and proteomic profiling. Although RNA and proteomic profiling data are often correlated, they may diverge due to post-transcriptional regulatory processes (Franks et al., 2017; Walley et al., 2016). Given that proteins generally are more directly involved in carrying out biological function than messenger RNAs, we hypothesized that a proteomic-based model would perform best. Additionally, combining both RNA and proteomic data was expected to produce the best-performing model by leveraging complementary information. To test this, we evaluated three model configurations: RNA-seq only features, proteomic profiling only features, and a combination model using both feature types.

For this test we used a comprehensive dataset from Walley et al. (2016), which contained paired RNA-seq (FPKM) and proteomic (dNSAF) measurements across 23 developmental stages and tissues in maize, enabling direct comparison between protein and RNA based models. We found that the RNA-only and protein-only models exhibited roughly equivalent performance, and combining the RNA and protein features did not improve model performance (*SI Appendix*, Fig. S2). This argued against our initial hypothesis that protein abundance measurements would be more directly predictive of pollen fitness. However, we were more surprised that modeling with both types of data gave no measurable boost to predictive performance, given that each dataset contained independent information that should be relevant to pollen fitness. For example, we expect a subset of transcripts in mature pollen to be poised for rapid translation upon pollen tube germination (Hafidh and Honys, 2021) (i.e., pollen fitness would be best predicted by RNA level for this subset), whereas more generally we expect protein level to be most predictive.

### 2.4 Measures of Pollen Specificity Contribute Heavily to Model Success

To better understand why mRNA and proteomic expression datasets did not differ markedly in information content relevant to pollen fitness, nor did they combine together to boost performance, we examined the relative predictive contribution of each feature across the RNA and protein expression models. Quantifying predictive contributions using the magnitude and sign of each feature’s weight provided an understanding of how strongly each feature contributed to positive (“strong fitness effect”) or negative (“no fitness effect”) predictions (*SI Appendix*, Fig. S3). We observed a recurring pattern in the proteomic expression features across many models: the weight of the expression value in mature pollen and that of the maximum level of protein expression across all samples tended to be similar in magnitude but oppositely signed, with max protein level always negative. This suggested that high expression in any tissue other than pollen tended to drive the model toward a classification of no fitness effect (*SI Appendix*, Fig. S3), and raised the hypothesis that a feature that explicitly captures pollen expression specificity would boost model performance.

To test this hypothesis, we constructed three measures of tissue specificity. The first feature, termed Is Pollen Max (IPM), is a binary variable indicating whether a gene model expressed maximally in pollen compared to all other measured tissue types. The second feature, termed Pollen Specificity Ratio (PSR), is a numeric variable representing the ratio of pollen expression to maximum non-pollen expression. The final feature, termed Non-Pollen Specificity ratio (NPS), is a numeric variable representing the highest ratio of expression in any non-pollen tissue relative to the maximum expression observed in all other non-pollen tissue types (*SI Appendix*, Fig. S4).

All three specificity features were calculated for all gene models in the dataset using both RNA-seq and proteomic profiling values (denoted with an R or P subscript, respectively), and incorporated into the modeling process (Fig. 3). PSR_*P*_ and IPM_*P*_ were among the most important predictors in both RNA and protein models, and were consistently assigned high positive weight relative to other features in the model (Fig. 3B & 3C). This indicated that models relied heavily on these features when predicting a fitness effect. Additionally, models that included these features showed a clear boost in predictive performance (Fig. 3A).

**Figure 3.**
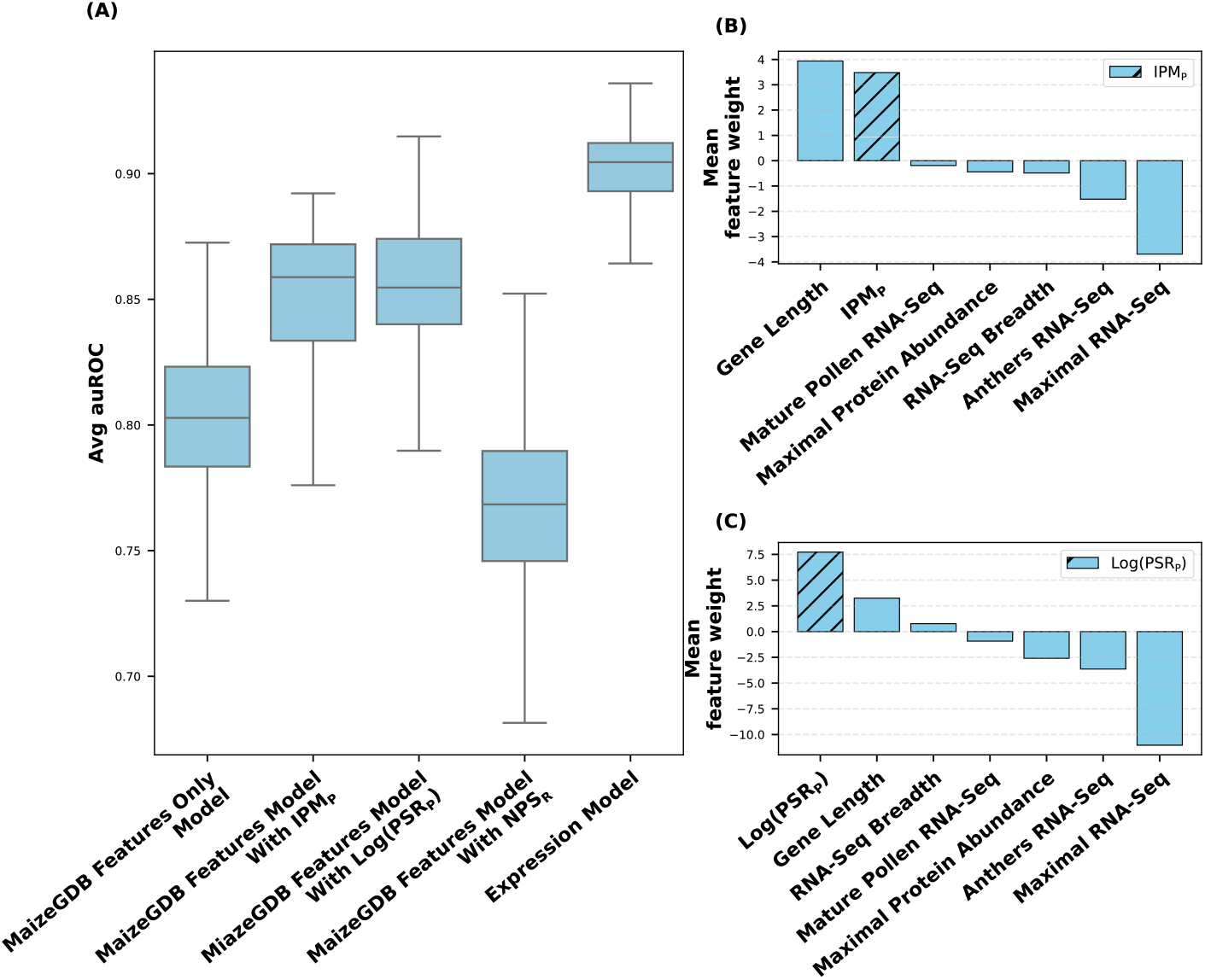
Derived expression specificity features substantially improve model performance. (*A*) Early expression-based models that included RNA and protein expression features (without specificity measures; *SI Appendix*, Table S2), achieved ~ 80% test-fold average auROC. Adding either protein abundance Is Pollen Max (IPM_*P*_) or protein abundance Pollen Specificity Ratio (PSR_*P*_) into the model increased performance by roughly 5%. The final Expression Model, which includes PSR_*P*_ and the RNA-seq based Non-Pollen Specificity ratio (NPS_*R*_) features, achieved an overall ~ 10% improvement in auROC. (*B,C*) Average feature weights across all 1,000 CV seeds from panel *A* show that IPM_*P*_ and PSR_*P*_ consistently received large positive coefficient weights, highlighting their strong influence on predicting fitness effect.

We next ran an automated exhaustive search over all unique combinations of expression and specificity-based features to identify a best expression feature subset. Specifically, our program tested all unique combinations of 14 different features (*SI Appendix*, Fig. S5A), training a separate model for each subset. This search yielded a best performing Expression Model with three features, two of which were specificity based: log-transformed Pollen Specificity Ratio for protein (positive weight, favoring strong fitness effect) and Non-Pollen Specificity Ratio for RNA-Seq (negative weight, favoring no effect). These two were also the two most frequently appearing specificity features among all of the top performing models (*SI Appendix*, Fig. S5A). In this search, both RNA-Seq and protein abundance derived versions of our three specificity features were included in order to explore which of these data types lead to best model performance. Notably, when visualizing the log-transformed PSR_*P*_ against transmission rates from the training set, a clear separation among pollen expression types becomes apparent: pollen minimal, pollen non-specific, and pollen specific (Fig. 4).

**Figure 4.**
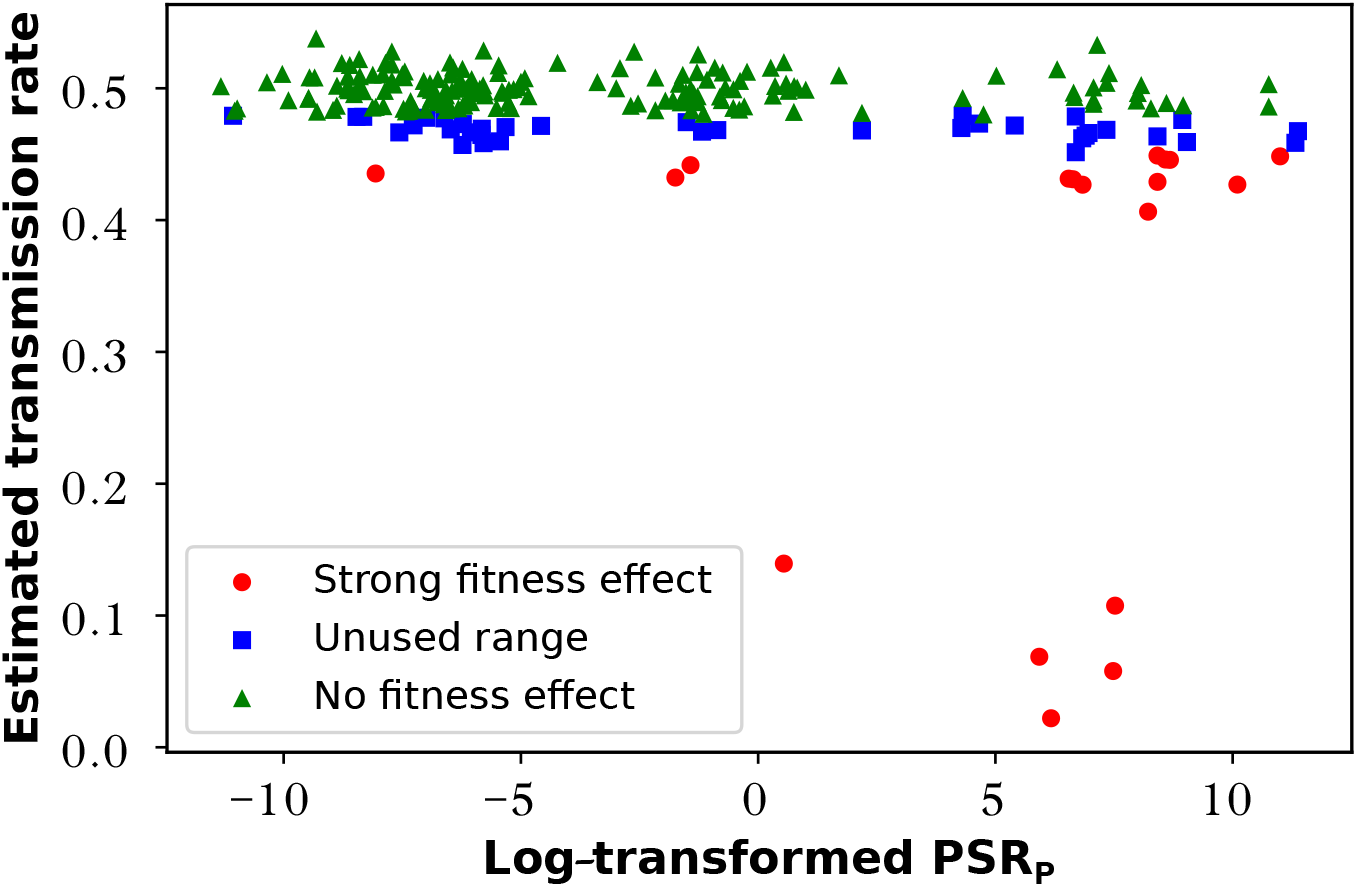
Gene models in the training dataset can be grouped based on pollen expression specificity. When estimated mutant transmission rate vs. the log-transformed Pollen Specificity Ratio (PSR_*P*_) is plotted, three distinct groups emerge based on expression pattern: pollen minimal (left), pollen non-specific (center), and pollen specific (right). Pollen minimal and non-pollen specific gene models dominate the no fitness effect class, whereas pollen specific gene models comprise a greater proportion of the strong fitness effect class.

These results help explain why no performance gain was observed when we combined RNA and protein expression features in a single model – the most important information is actually pollen specificity, which was not being fully captured without our derived features. Therefore, we incorporated these specificity features into all subsequent model formulations with expression data.

### 2.5 Additional Genomic Features Can Increase Model Performance

We also explored whether the large collection of genome sequence-derived and epigenomic features available from MaizeGDB (Sen et al., 2023) and Hufford et al. (2021) could provide predictive power distinct from gene expression data. Specifically, this experiment had two goals: first, to develop a model that uses only genome sequence derived features; and second, to determine whether including such features could boost the performance of expression-based models.

Due to the large number of features (497 features), these features were analyzed in subsets by category, for example, nucleotide dimers and protein structure (*SI Appendix*, Table S3). A systematic process using four different models was applied to each subset to assess predictive capability in a feature reduction screen. (1) The entire feature subset was used to build a model, (2) the feature subset was down-sampled based on a feature-to-outcome correlation threshold, (3) the full feature subset was combined with the best-performing expression model, and (4) a reduced feature model was built by selecting top weighted features across the three previous models to use in combination with the best expression model. Models 1 and 2 evaluated the predictive power of genomic sequence features alone, whereas models 3 and 4 assessed their ability to enhance expression based models (see Methods Section 3.5 for a detailed description).

For each subset, the auROC outcome values of the four models were compared using three evaluation measures that we created: “Baseline Predictive Power”, “Down-sampling Gain”, and “Expression Boost”. Baseline Predictive Power was calculated as the maximum of the difference in auROC between each model (model 1 and model 2) relative to a 65% auROC baseline. This helped to assess whether the feature subset alone had meaningful predictive power. Down-sampling Gain was calculated as the difference in auROC between model 1 and model 2. This helped assess whether removing low-correlation features could improve the predictive signal of the feature set. Expression Boost was calculated as the greater of the two differences in auROC of model 3 and model 4 relative to the best expression model auROC. This helped to assess whether the genomic feature subset could improve the performance of expression-based features.

Each feature set was scored as having high, moderate, or low predictive potential based on our evaluation measures (*SI Appendix*, Table S3). This identified 16 composite feature conditions, used in selecting the most information-rich features across all the genomic subsets (*SI Appendix*, Table S4). A final round of modeling incorporating these conditions resulted in streamlined feature subsets that combined to generate the highest performing Genome Model (auROC of 85.64%) and Combination Model (auROC of 92.03%) (Fig. 2, *SI Appendix*, Table S1). The Genome Model, based on specific nucleotide dimer, codon usage, exon length and protein composition features, did achieve a strong auROC value, but overall was the weakest of the three final models. In contrast, the Combination Model outperformed both the Genome Model and Expression Model, using PSR_*P*_ along with specific protein composition, codon usage, and evolution-relevant features (e.g., singleton status in the maize genome). This revealed that certain genomic features add independent information content that complements the PSR_*P*_ measure of expression specificity, improving predictive capability.

### 2.6 Genome Scale Application of Pollen Fitness Models Suggests Expression Driven Models Can Generalize as a Predictive Tool

To assess the generalizability of the final three models (Fig. 2), we performed two analyses. In the first analysis, the Genome, Expression, and Combination models were used to predict fitness effects for the B73v5 gene models that had all of the necessary feature data (*SI Appendix*, Table S1) from our sources (e.g. MaizeGDB). After excluding gene models contained in our training set, this produced a set of approximately 13,800 gene models. We then visualized the relationship between pollen specificity (Log(PSR_*P*_)) and predicted fitness effect for each model (Fig. 5). As expected, these distributions showed that the two models incorporating expression features tended to assign the strong fitness effect class to pollen specific gene models. In contrast, the Genome Model assigned the strong fitness effect class across the entire range of PSR_*P*_ values, suggesting that the genomic features used are likely to represent aspects of the training set that are distinct from pollen specific expression. Additionally, the Genome Model produced a much higher number of strong fitness effect predictions than either model incorporation expression features (Fig. 5D).

**Figure 5.**
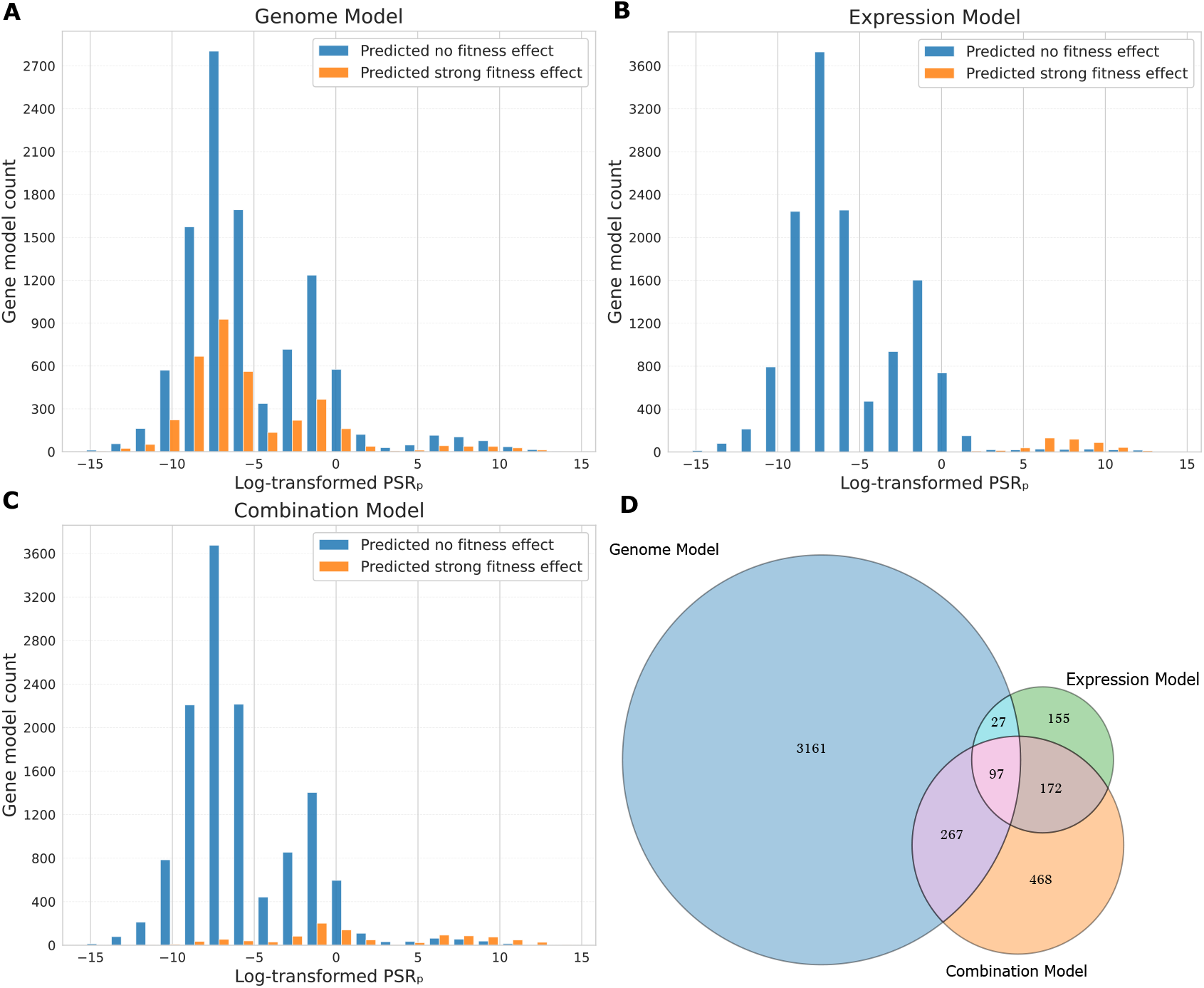
Models incorporating expression specificity features appear to generalize well to genome scale prediction. *(A-C)* All three model types were applied to predict fitness class across ~13,800 gene models. Predicted fitness classes, strong and no effect, are shown with respect to their log-transformed Pollen Specificity Ratio PSR_*P*_. *(A)* The Genome Model predicted the largest number of strong fitness effect gene models, most of which were non-pollen specific. *(B)* The Expression Model predicted the fewest strong fitness effect gene models, primarily within the pollen specific range of PSR_*P*_. *(C)* The Combination Model produced an intermediate number of strong fitness effect gene model predictions, capturing gene models of all three expression groups (pollen minimal, pollen non-specific, pollen specific). *(D)* The Venn diagram shows the overlap and distinctions among strong fitness effect predictions from the three different models. Total strong fitness effect predictions: Genome = 3552, Expression = 451, Combination = 1004

In our second analysis exploring the final models’ predictive capability, we analyzed an independent set of transposon insertions from the maize BonnMu population (Marcon et al., 2020; Win et al., 2024). This large population, designed for functional genomic studies, includes 425,000 genomically mapped heritable mutations covering over 80% of maize gene models. Because the mutagenized DNA in this population was generated by transmission solely through the pollen, insertions in genes important to pollen fitness would be less likely to be recovered as heritable, and therefore less likely to be present in the BonnMu dataset. By contrast, gene models that have no effect on pollen fitness would not be associated with reduced recovery. To test whether our three final models met this expectation, we matched the predicted classes (strong fitness effect vs. no fitness effect) from each to the gene models represented in the BonnMu dataset. The number of BonnMu insertions per gene model, categorized by genomic section (e.g., exon, intron, etc.), were then counted, and the mean differences between the two predicted fitness classes were compared (Fig. 6).

**Figure 6.**
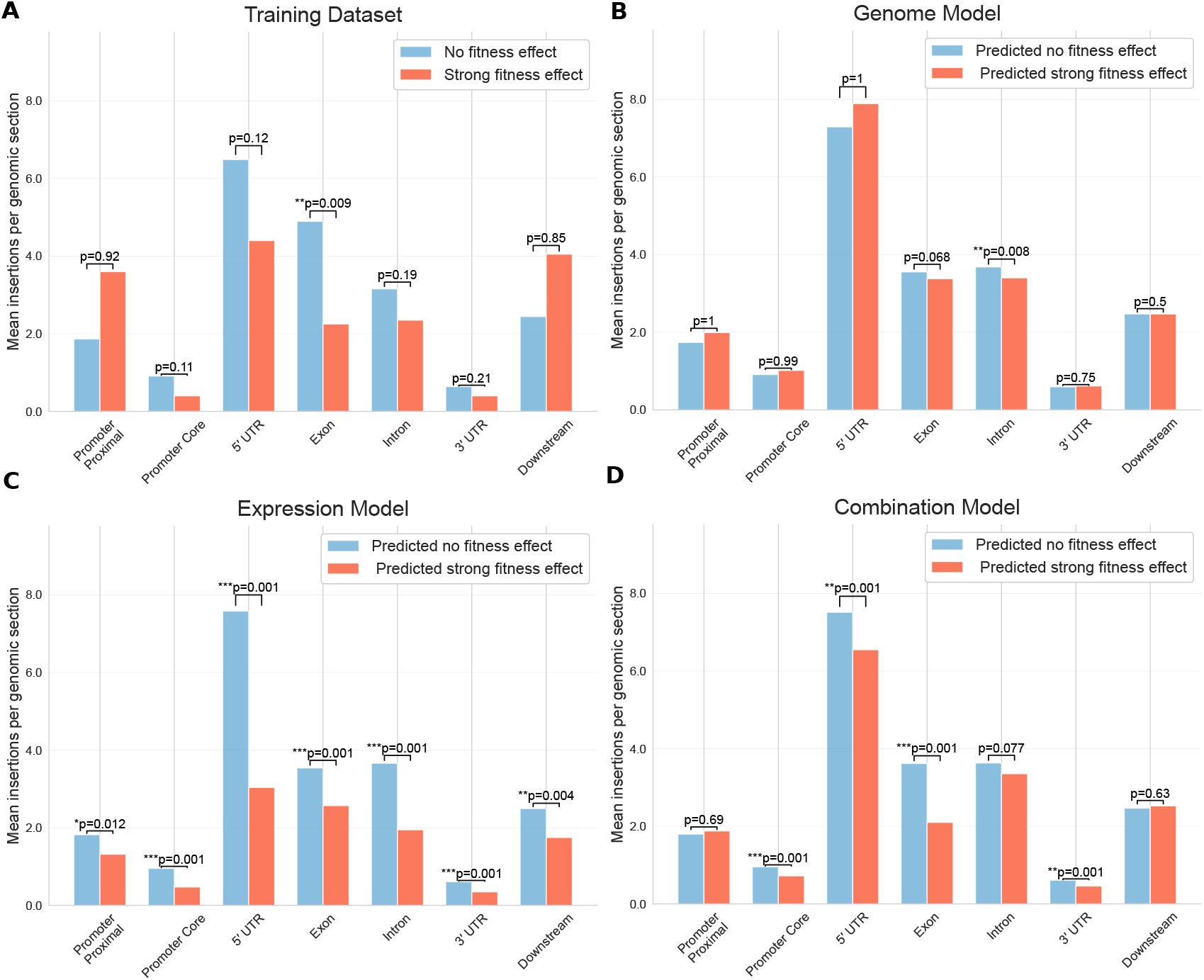
The Expression and Combination Models successfully predict expected relative frequencies of heritable insertions in the BonnMu transposable element population across the maize genome. *(A-D)* Mean BonnMu insertion count per genomic section in comparator gene model sets, grouped by class. Expectations are that genes important for pollen fitness would be underrepresented among heritable BonnMu insertions, relative to genes with no effect on pollen fitness, because mutations disrupting such genes are less likely to be transmitted through the pollen and recovered as heritable. Statistical values were obtained by 1,000 label shuffing permutations of gene model and class prediction labels. Reported p-values correspond to one-tailed tests (strong fitness effect counts < no fitness effect counts). *(A)* As a control, the training dataset shows the expected trends, with exons exhibiting a significant insertion count difference. *(B-D)* Although the Genome Model predictions did not show the expected trends, both the Expression Model and Combination Model showed significantly reduced frequencies in predicted strong fitness vs. no fitness effect classes across at least four of the seven genomic sections. Proximal Promoter = −2100 to −100 bp from the transcription start site; Promoter Core = −100 bp to the transcription start site; Downstream = +1 to +2100 bp downstream of transcript end.

The BonnMu dataset maps insertion locations in the population to seven distinct genomic sections: Promoter Proximal, Promoter Core, Five Prime UTR, Exon, Intron, Three Prime UTR, and Downstream. We expected to observe a difference in average counts for all of the genomic sections except Promoter Proximal and Downstream. These two sections span thousands of basepairs upstream and downstream of the annotated gene models, and thus mutations that influence gene function would be uncommon (i.e., insertion into a critical regulatory site). As a control, we assessed the distribution of insertions in gene models from the training set, based on our class assignment of strong fitness effect versus no fitness effect (Fig. 6A). The results confirmed our expectations: insertional counts average lower for the strong fitness effect gene models in every genomic section, except for promoter proximal and downstream.

To evaluate whether the observed differences in average BonnMu insertion counts in these analyses could have occurred by chance, permutation tests were performed (Methods 3.7.2). The resulting p-values helped to assess whether the test set results were significantly different than would be expected by chance. Using this approach, we found that the Genome Model (Fig. 6B) showed a significant difference only in the Intron section. In contrast, the Expression Model (Fig. 6C) exhibited significant differences across all genomic sections. Similarly, the Combination Model (Fig. 6D) showed significance in all sections except Promoter Proximal, Intron, and Downstream regions. Notably, in line with the training set, the Combination Model exhibits the strongest signal in the Exon section.

A potential confounding factor for this analysis is that transposable element insertion rates are non-uniform across the genome. In fact, *Mutator* elements are known to preferentially insert near transcription start sites, with data also pointing towards increased insertion rates at highly expressed genes (Zhang et al., 2020). Therefore, genes with pollen specific expression could have reduced accessibility to insertion. However, the proportional differences in the representation of BonnMu insertions in the different genomic sections are difficult to explain by generalized reduced accessibility of genic regions of pollen specific genes. This is especially notable in the Exon vs Intron insertion frequencies based on predictions of the Combination Model (Fig. 6D). Thus the data suggest that pollen fitness effects are very likely to be a significant driver of the observed insertion rate differences.

Finally, we examined whether higher probability predictions of a strong fitness effect were associated with a more severe or a more likely fitness effect. To explore this idea, we took the predicted strong fitness effect genes from each of the three models and divided them into upper and lower halves, based on predicted probability value of having a strong fitness effect. If high probability predictions were associated with a more severe or more likely fitness effect, then we would expect that the gene models associated with the “upper half” group would exhibit a lower BonnMu insertion count, i.e., a lower insertion recovery rate in the BonnMu population, than the paired “lower half” group. For the Genome Model (*SI Appendix*, Fig. S6A), all genomic sections except Intron exhibited this behavior, with a significant signal in Exon. For the Expression Model (*SI Appendix*, Fig. S6B), all genomic sections except Promoter Proximal and Intron exhibited this behavior. For the Combination Model (*SI Appendix*, Fig. S6C), all genomic sections except Promoter Proximal exhibited this behavior. Notably, the Expression Model and Combination Model showed this trend significantly in the Five Prime UTR, in agreement with our other analyses (Figures 5 & 6) that these two model types appear to be the most generalizable across the genome. These results generally support the idea that the fitness outcome probabilities assigned by all of our predictive model types, at least to some degree, are encoding severity as well as likelihood of fitness effect.

## 3 Discussion

The availability of genome-scale datasets is expanding rapidly across the Tree of Life as data-generation costs decrease; however, the exact relevance of each of these data types to a gene’s ability to influence phenotype (i.e., biological function) is unclear. Here, we developed three ML models to predict gene-specific quantitative phenotypes from genotype-linked omics data. Our models served not only as predictive tools, but also as a means to assess the relative contribution of each genome-scale data type to predicting phenotype.

Our Genome Model achieved the lowest auROC, likely reflecting the limited training dataset size. While this model could predict phenotype from sequence derived features alone, our followup analyses indicate that it generalized across the genome the least well. However, the model’s ability to achieve an auROC of 85% suggests that both generalization and performance could improve with additional data. Increased training set size and additionally informative genomic features would clearly benefit the model. For example, features derived directly from genome sequence that encompass information about a gene’s pollen specificity may be able to improve performance by providing another avenue to the specificity information that the expression data currently provides in the other two models. Our Expression Model (auROC 90%) and Combination Model (auROC 92%) both achieved very strong performance, substantially higher than the Genome Model (auROC 85%). Notably, the Combination Model also generalized best across the genome, as evidenced by the BonnMu analysis. In particular, not only did the Combination Model predict that a reasonably expected number of gene models would show a strong fitness effect on mutation, but these gene models also showed the strongest BonnMu insertion differential in Exon. This indicates that expression features and genomic sequence features can act synergistically when predicting phenotype. Both the Expression and Combination Models incorporated measures of expression specificity, highlighting its unique importance as a predictive feature.

Specifically, we engineered three features to quantify expression specificity. The features Is Pollen Max (IPM) and Pollen Specificity Ratio (PSR) quantify pollen expression specificity, whereas Non-Pollen Specificity ratio (NPS) quantifies non-pollen expression specificity. Interestingly, pollen specificity using proteomic data gave better predictive performance than using the analogous RNA-Seq data, whereas the opposite held for quantifying non-pollen specificity. These differences may be attributable to post-transcriptional regulatory processes in pollen tissue. Across many angiosperms, differences between transcript level and protein abundance in pollen have been observed (Hafidh and Honys, 2021; Walsh et al., 2020; Hafidh et al., 2018; Robinson et al., 2021). Proteomic data from tomato pollen has raised the idea that accumulation of certain proteins prior to pollination primes the system for the subsequent developmental stage (Chaturvedi et al., 2016). This enrichment of proteins required for rapid function (e.g., pollen tube growth) may make protein abundance data a better marker of pollen specificity than transcript count for the same genes. Because pollen specificity quantification was a key driver of predictive performance, a more relevant signal of pollen specificity likely improved our models. Broadly, protein and transcript derived specificity metrics may capture distinct regulatory layers, explaining the differing utility across specificity quantification.

The central importance of pollen specificity to the predictive success of our models could stem from the somewhat extreme nature of maize pollen. A single maize plant generates millions of pollen grains (Uribelarrea et al., 2002), and the floral structure (silk) on which they land has the potential to capture hundreds of these grains, only one of which will outcompete the others to successfully fertilize the embryo sac to generate a single seed (kernel). This competition requires pollen tube growth, and, in comparison to other plants, maize pollen tube growth rates are among the fastest known, and the distances traveled are among the longest (Zhou et al., 2017; Bedinger and Fowler, 2009). To meet these demands, selection may have favored the evolution of highly specialized genes whose expression is limited to pollen, conferring these extreme biological characteristics. Consequently, pollen expression specificity would have emerged as a key driver of model performance, marking genes that are specialized contributors to maize pollen fitness. Further research is needed to determine whether expression specificity is a predictive proxy for phenotype that can be generalized to other developmental systems and other species.

The discovery of pollen specificity as the major driver of pollen fitness prediction in our models raises two additional points worth considering. First, an open question: why are there a large number of non-specific genes expressed in pollen despite the energetic costs? In our training dataset, pollen non-specific and specific genes are associated with similar average protein abundances, suggesting that the energetic burden of gene and protein expression is similar for both groups. However, our study suggests that pollen non-specific genes contribute far less to fitness than pollen specific genes. One speculative explanation is that mutations that reduce or eliminate the transcription of broadly expressed, non-specific genes only in pollen (i.e., a very specific kind of regulatory mutation) are very rare. Alternatively, pollen non-specific genes may contribute to fitness at levels generally below the detection limits of the EarVision system, but enough to provide a selective advantage that maintains expression in pollen. Given the inherent limitations of our small training dataset, it is also possible, but unlikely in our view, that the dataset systemically underrepresents particular types of strong fitness effect genes, limiting the generalizability of model results (e.g., if low abundance proteins not represented in available proteomics data are also more generally pollen non-specific).

Second, the possibility that pollen fitness is largely conferred by genes expressed in a pollen specific manner is relevant to a long-standing hypothesis in plant evolutionary genetics that the gametopyhtic phase plays a role in purging plant genomes of deleterious mutations (Beaudry et al., 2020; Nelms and Walbot, 2022; Immler and Otto, 2018). In this view, the haploid gametophytic phase serves as a “genetic filter” and helps eliminate deleterious mutations that could influence the sporophyte (i.e., the most visible, diploid phase of the plant life cycle). Mutations reducing or eliminating gene function, usually recessive in diploids, can accumulate in sporophytically-expressed genes in a population, remaining phenotypically masked in heterozygotes. However, because pollen is haploid, selection on the gametophyte is expected to remove such alleles, if the corresponding genes are also expressed in pollen (i.e., such mutations would reduce pollen fitness). Our models’ association of genes with pollen non-specific expression (i.e., those with potential to influence sporophytic phenotypes) with a lack of strong pollen fitness phenotypes suggests that gametophytic purging of such alleles is more limited than expected, at least in maize. A more extensive pollen fitness dataset would enable a more definitive evaluation of the extent of the hypothesized gametophytic purging effect on sporophytically-expressed genes.

In summary, this study presents a framework for using omics-scale data to associate single genes with a specific phenotype. Using interpretable ML methods, it is possible not only to build successful models, but also to efficiently evaluate a large number of omics-scale datatypes to identify high-value features. As high-throughput phenotyping data become more available, similar pipelines could be applied to develop genotype to phenotype models in other organisms, e.g., for seedling phenotypes of the model plant Arabidopsis (Wan et al., 2025). Such studies should help evaluate whether expression specificity is a general marker for phenotypic influence across different tissue types and species.

## Mthods

### 3.1 Ds-GFP Pollen Fitness Dataset

*Ds-GFP* lines were cultivated at the Oregon State University Botany and Plant Pathology Farm in Corvallis, OR as described (Ruggiero et al., 2025). Briefly, lines containing insertions into targeted gene models were requested from the Maize Genetics Cooperation Stock Center. All insertion alleles were validated via PCR followed by Sanger sequencing, using a custom primer design pipeline, to confirm insertion coordinates (*SI Appendix*, Data S1). For generating experimental crosses with specific *Ds-GFP* alleles, GFP-positive kernels from outcrosses to the W22 *r c1* inbred line (i.e., heterozygous plants) were grown to maturity, sampled for DNA, and genotyped using allele-specific primers paired with primers internal to the *Ds-GFP* element (*SI Appendix*, Data S1). These GFP-positive heterozygotes were used as parents for controlled, competitive pollinations to a *c1* homozygous tester line, using fresh pollen and extensive silk area on recipient ears. To obtain transmission rates, harvested ears were imaged with the Maize Ear Scanner.v2 and analyzed via the computer vision pipeline EarVision.v2; ears that did not satisfy established quality thresholds (Ruggiero et al., 2025) were hand annotated in ImageJ. Raw GFP and non-GFP kernel counts were analyzed via a GLM-based pipeline in R to obtain estimated transmission rates and associated p-values for each allele.

### 3.2 Ds-GFP Gene Model Feature Association

All *Ds-GFP* insertional elements were mapped relative to the B73v5 maize genome annotation (Woodhouse et al., 2021a; Hufford et al., 2021) using the Python library PyBedTools (Dale et al., 2011) to identify insertion locations in B73v5 gene models. The preprocessing pipeline also included project specific data-wrangling steps to resolve issues such as allele naming inconsistencies and removal of multiple gene model associations, resulting in a *Ds-GFP* dataset with 305 unique gene model associations.

For each gene model, we retrieved features from MaizeGDB FeatureBase (Sen et al., 2023). MaizeGDB organizes available features into broad categories: Sequence Features, Gene Structure, Expression, Chromatin Features, Count, Correlation, Varionomic, and Miscellaneous. Within each category, subsets of features were chosen based on their potential relevance to pollen function. In addition to these, information related to gene model singleton or paired status relative to the ancient whole genome duplication of maize, and each gene model’s location in the two maize subgenomes (M1 and M2), was extracted from a recent analysis of the maize pangenome (Hufford et al., 2021) (*SI Appendix*, Data S6). This process ultimately yielded over 600 features with potential relevance to pollen fitness.

### 3.3 Constructing Pollen Specificity Features

After identifying the importance of pollen specificity, we derived multiple measures of tissue specificity that we hypothesized could provide additional predictive information. Each feature was calculated across the 23 tissues assayed in a paired RNA expression and protein abundance dataset (Walley et al., 2016).

We used the following matrix formulation: let *X* ∈ ℝ^*n*×*m*^ denote the RNA expression matrix or protein abundance matrix, where n is the number of gene models and m is the number of tissues. Let *i* be index of gene model (*i* ∈ {1, …, *n*}), *j* be the index of tissue (*j* ∈ {1, …, *m*}). Let *T*_*np*_ be the set of non-pollen tissue indices (*T*_*np*_ ⊂ {1, …, *m*}). In the following, *ϵ* is a fixed small numerical value to avoid division by zero.

#### 3.3.1 Is Pollen Max (IPM)

This binary feature (true/false) denotes whether expression for a given gene model was maximal in pollen tissue compared to all other tissues:

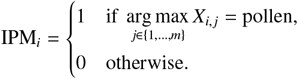

#### 3.3.2 Pollen Specificity Ratio (PSR)

This numeric quantity represents the ratio of pollen to non-pollen expression for a given gene model. The numerator is the pollen expression value and the denominator is the maximum non-pollen tissue expression value:

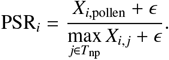

#### 3.3.3 Non-Pollen Specificity ratio (NPS)

This numeric quantity represents the non pollen tissue-specificity of maximum expression value for a given gene model. For each gene model, the numerator is the highest non-pollen tissue expression value, and the denominator is the second-highest highest non-pollen expression value:

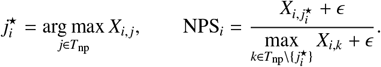

During experimentation the three specificity features were calculated using both RNA-Seq and proteomic profiling data, denoted by an R or P subscript (e.g. PSR_*P*_). We found that PSR provided greater predictive power in combination with other features when calculated from proteomic profiling data. Conversely, we found that NPS and IPM were more predictive when calculated with RNA-Seq data as opposed to proteomic profiling data (*SI Appendix*, Fig. S5A). Additionally, log transformed PSR_*P*_ further improved model performance metrics compared to using the non transformed version.

### 3.4 Machine Learning Pipeline

All analyses were implemented in Python version 3.10.12 using standard scientific and machine learning libraries (pandas, scikit-learn, NumPy, Matplotlib and Seaborn). We provide the analysis code pipeline as described in *SI Appendix*, Code S1.

#### 3.4.1 Data Preparation

The input dataset consisted of rows corresponding to B73 V5 gene model IDs, columns corresponding to features, and the outcome variable (transmission rate). The continuous outcome variable was discretized into a binary fitness class. Gene models with a transmission rate >48% were assigned to the no fitness class (labeled as 0), while gene models with transmission rate <45% were were assigned to the strong fitness effect class (labeled as 1). Categorical features (e.g., subcellular localization and paired/singleton status (Hufford et al., 2021)) were one-hot encoded prior to modeling. For each model run, any gene model that contained a missing value (NaN) in a selected feature column was excluded.

In this dataset the number of input features (600+) is very large compared to the number of data points (~227). This creates an extremely large feature space that the machine learning algorithm must search in order to identify meaningful combinations of features that can successfully predict outcome. Feature selection is necessary in this situation to enable the potential of machine learning algorithms to identify a successful predictive solution – that is, to enable an ML model to identify an equation and its parameters that successfully predicts outcome from features for every input datapoint. We constructed many different feature subsets, using biological reasoning and information gleaned from the modeling process to guide the search for features that were potentially meaningful in predicting fitness outcome. In total, over 350 feature subsets were tested, using the procedures described below.

#### 3.4.2 Modeling Approach

The primary model type used was logistic regression with Lasso (L1) regularization. Models were implemented using the Python Library scikit-learn (version 1.7.0). Random forest models were also evaluated but did not provide a significant performance improvement, so we used logistic regression for analyses described in the project. Two hyperparameters were tuned during model training for each feature subset explored: strength of regularization and the class-weighting parameter. Class-weighting was implemented to address the class imbalance present in our dataset; there were ~ ten times as many “no fitness effect” gene models as “strong fitness effect” gene models.

#### 3.4.3 Cross-validation

To train and evaluate each model, three-fold cross-validation was used. Stratification was applied to ensure class balance within each partition, using the scikit-learn StratifiedkFold function. For every feature subset, the model was trained on two of the three partitions and evaluated on the unseen third partition. This was repeated twice more to ensure each partition served as an evaluation set once. Because our dataset was relatively small for training machine learning models, cross-validation served two important purposes: it maximized the use of available data for model training and provided an estimate of model overfitting.

For initial model analysis, three-fold cross-validation partitioning was performed using a single random seed. For final performance reporting, models were evaluated as the average across 1,000 random seeds (0-999). This approach provided a more robust estimate of model performance.

#### 3.4.4 Model Evaluation Metrics

Model performance was evaluated using multiple metrics: auROC, as well as sensitivity, specificity, and accuracy at the Youden’s index threshold. auROC was the primary metric for overall performance evaluation because it is well suited for binary classification and for models that output class prediction probability estimates where ability to achieve a balance between strong sensitivity and strong specificity is important. auROC evaluates the trade-off between sensitivity (true positive rate) and specificity (1 - false positive rate) across different prediction probability cutoffs (e.g. strong fitness effect defined as an outcome of *p* >.5813). This allowed us to choose a probability cutoff using Youden’s index, which is the cutoff that provides the “best” balance between sensitivity and specificity, in the sense that the Youden’s index performance outcome is the closest one can get to achieving both 100% specificity and 100% sensitivity at the same time. Model metrics were reported as the mean across all three test partitions in cross-validation, and also examined for each individual partition to monitor potential overfitting.

#### 3.4.5 Feature Interpretation

A key benefit of logistic regression is the linear structure by which features are combined through weighted coefficients and the interpretability that this brings to the modeling process. The logistic regression equation is defined as:

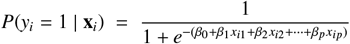

where *P*(*y*_*i*_ = 1 | **x**_*i*_) is the probability that gene model *i* has a strong fitness effect outcome, given its feature vector **x**_*i*_ = (*x*_1_, …, *x*_*ip*_), scaled by corresponding feature weights *β*_1_, …, *β*_*p*_. Because the predicted probability value is based on a linear combination of features and their weights, we examined the learned feature coefficients to assess each feature’s contribution to predicting strong fitness effect vs. no fitness effect. For all non-negatively valued features, positive coefficients increase the probability of a strong fitness effect prediction (*P* → 1) by increasing the magnitude of the linear combination of features and weights in the positive direction (towards class 1 - the strong fitness effect class), while negative coefficients decreases the probability (*P* → 0) toward class 0 which is the no fitness effect class. All features in our model are non-negative except for log-transformed PSR_*P*_, whose coefficient has its own unique interpretation. All feature values were Min-Max scaled to the same range (0-1), enabling interpretive comparison across features that have very different scales. Examining the magnitude and sign of feature weights provides an interpretable measure of feature importance that guided our modeling process. Additionally, L1 regularization “shrinks” the coefficients of features that are less informative for successful prediction towards zero. This has the effect of encouraging successful models with fewer more highly weighted (therefore potentially explanatory) features, as opposed to models that use many low weighted features. Therefore, L1 regularization served the dual purpose in our modeling process of performing automated feature selection while seeking the simplest possible well performing model.

### 3.5 Genome Sequence Derived Feature Set Evaluation

To assess whether genome sequence-derived features could independently produce a successful “genome only model” and improve an expression-based model, we evaluated the large set of genomic features (Sen et al., 2023; Hufford et al., 2021) in smaller groupings. These subsets were organized following the MaizeGDB categorization (e.g. tri-mer counts), resulting in 15 distinct feature subsets. The complete list of subsets is provided in table (*SI Appendix*, Table S3).

#### 3.5.1 Experimental design

To assess the predictive power of each feature subset, we constructed four different models for evaluation. Model 1 (genome-only): includes all of the features in the subset. Model 2 (downsampled genome-only): includes a downsampled version of the feature subset (see Section 3.5.2). Model 3 (expression + genome): includes the entire feature subset plus the features associated with the best performing expression model at the time of experimentation (log(PSR_*P*_), IPM_*P*_, gene length, gene breadth, mature pollen RNA expression, anthers R1 RNA expression, max protein abundance, max RNA expression). Model 4 (expression + hand-curated genome): includes the best performing expression model with a hand-curated collection of the feature subset, selected based on insights from Models 1-3. Using the results of Models 1-4, we assessed the predictive potential of the feature subsets using custom measures explained in Section 3.5.3.

#### 3.5.2 Downsampling procedure

For Model 2, downsampling was necessary because the number of genomic features (~500) was large compared to the number of gene models (~150), preventing the L1 regularization from identifying high information content features. Therefore, we implemented a preliminary feature selection process to identify a reduced set of potentially meaningful features. Model 2 downsampling was performed by selecting the features most highly correlated with outcome. For continuous numeric features, correlations were calculated using the point-biserial correlation coefficient, a variant of Pearson’s correlation coefficient that is appropriate for comparing a vector of continuous numerical values with a vector of binary values. For categorical features, correlations were calculated using the Phi coefficients, a correlation coefficient that is appropriate for comparing two vectors of binary values. The exact correlation cuttoff was evaluated for each feature subset. The top ~ 25% of features with strongest correlations were retained for Model 2 evaluation, with the goal of reducing each subset to less than ~50 features.

#### 3.5.3 Evaluation Measures

We constructed three evaluation measures: *Baseline Predictive Power, Downsampling Gain*, and *Expression Boost*. The purpose of Baseline Predictive Power and Downsampling Gain was to evaluate the predictive potential of genome only models, whereas the purpose of Expression Boost was to assess the extent to which genomic features could enhance an expression based model. Every feature subset was evaluated using these three measures.

##### Baseline Predictive Power

To calculate Baseline Predictive Power, we first defined a minimum auROC of 65%. We then compared this baseline to the average cross-validation test-fold auROC of Models 1 and 2. The maximum difference between each model’s auROC and 65% was assigned as the Baseline Predictive Power for the feature subset.

##### Downsampling Gain

Downsampling Gain was calculated as the difference between the average cross-validation test-fold auROC of Model 1 and Model 2.

##### Expression Boost

To calculate Expression Boost, we first identified the best expression-based model auROC as 87% (best expression model auROC at time of experimentation). We then compared this auROC benchmark to the average cross-validation test-fold auROC for Models 3 and 4. The maximum improvement of either Model 3 or Model 4 over the benchmark was assigned as the Expression Boost value.

After calculating the three measures for each feature subset, we developed a scoring system for predictive potential. For each measure, we plotted the distribution of per model measure values across all feature subsets to visualize the distribution’s modes and spread. Based on these distribution properties, custom performance ranges were defined and assigned scoring labels of low, moderate, or high predictive potential (*SI Appendix*, Table S3). Each feature subset was assigned a label for each measure according to where its measure value fell within these performance ranges.

#### 3.5.4 Composite Feature Conditions

After all feature subsets had been labeled, composite feature conditions were defined. These conditions specified rules for building new feature subsets of manageable size (e.g., include only subsets labeled as high predictive potential for Baseline Predictive Power and Downsampling Gain) across all categories, to be used in the final round of genomic feature modeling. A full list of composite feature conditions is provided in *SI Appendix*, Table S4. The Genome Model and Combination Model were derived from these composite feature conditions.

### 3.6 Feature Subset Search

We developed a Python script to evaluate the model outcome (auROC) for every feature combination in a given feature set. Because the number of combinations increases exponentially with feature count, the script was applied only to small feature sets that had already been identified as meaningful in prior experimentation. These include the collection of high interest expression features in *SI Appendix*, Fig. S5 and the best performing composite condition feature sets from *SI Appendix*, Table S4. The Genome Model, Expression Model, and Combination Model were all identified using this script to select the final feature set.

### 3.7 Genome-Scale Application

Recognizing that our model training process always used cross-validation test sets due to the very limited number of gene models with fitness outcomes for training, and that several parameters were tuned using these cross-validation test sets without a truly independent held-out test set, we decided to perform additional evaluation of the final models using the entire maize genome, excluding gene models from our training dataset. We assembled a set of ~13,800 B73 V5 gene models that had precomputed MaizeGDB features, and then applied our Genome Model, Expression Model, and Combination Model to predict fitness effect probability for each of these gene models. For final class prediction probability cutoff we used the average Youden’s index across all test partitions from the cross-validation seed test explained in Section 3.4.3.

#### 3.7.1 Assessment of Model Predictions via Analysis of BonnMu Population Insertion Frequency

The BonnMu population dataset (Win et al., 2024) annotates the set of maize gene models with counts of presumed heritable transposon insertions in each gene ‘section’ (e.g., CDS, intron). BonnMu insertion counts into each genomic section were extracted for the ~13,800 gene models used for genome-wide application of the final models. These were then divided into two groups based on the class (strong fitness effect or no fitness effect) assigned by each final model. For each genomic section, for each gene model, the insertion counts in that section of the gene were summed and divided by the number of genes in that predicted class group (strong fitness effect, no fitness effect). This computes the average BonnMu insertion count per genomic section for each predicted class group. The per-genomic section difference between class groups was used as an indicator of model performance. The same procedure was applied to the *Ds-GFP* training dataset, using the class labels assigned based on transmission rate to define the two groups.

For each of the three final models, we further analyzed the gene subsets predicted to have a strong fitness effect. Gene models were ranked by predicted probability value, divided into upper and lower 50% groups by predicted probability, and then analyzed to determine the BonnMu insertion frequencies as described above for each group.

#### 3.7.2 Permutation test

For each BonnMu insertion frequency analysis, a permutation test was used to assess statistically the differences in insertion count average between the two classes for each genomic section. Specifically, we tested the null hypothesis that predicted class labels were unrelated to the average insertion counts:

*H*_0_: The class-wise averages are equal.

*H*_*A*_: The strong fitness effect class average is strictly less than the no fitness effect class average.

Across all gene models with class assignments, the predicted class labels were randomly permuted 1,000 times (preserving class size), using NumPy’s permutation function with a fixed random seed (42) for reproducibility. For each permutation, we computed BonnMu insertion frequencies and recorded the average count difference for each genomic section. This generated a null distribution of average count differences, i.e., when there is no association between predicted class label and average BonnMu insertion count. One-sided p-values were reported as the proportion of average count differences at least as extreme as our observed average count differences.

### 3.8 Data Availability Statement

Data and code supporting the findings of this study are available in the *SI Appendix*. Sanger sequencing data for Ds-GFP insertion flanking sequences are available in Genbank, Accession numbers PX761006 - PX761265.

## Supporting information

AlleleGenomicInformation

AlleleKernelCounts

TransmissionRateData

GeneModelsWithMultipleAlleles

FullInputDataset

Huffordetal.SyntenyDataset

ModelTrainingDatasets

GenomeWideDataset

GenomeWideDatasetPredictions

BonnMuDataset

ProjectCode

Supporting_Information_and_Supplemental_File_Index

## Author Contributions

JF, MM: designed the study. JF, ZV: designed and implemented data collection procedures. SM, MM: designed and implemented the modeling process. SM: prepared figures and wrote the manuscript. MM, JF: edited the manuscript. ZV: read and evaluated the manuscript.

## Acknowledgments

Marilyn Leary provided bioinformatic support on the phenotyping pipeline, including writing code for automated allele-specific PCR primer design. H. Bell, H. Flieg, K. Kress, T. Lawrence, J. McDonald, G. Michna, I. Nicholson, A. Perez, C. Waite, and H. Woodard provided invaluable assistance for dataset generation on the project in lab and field. Special thanks to Hugo Dooner and Charles Du for generating the *Ds-GFP* insertion population, as well as providing recommendations and suggestions for effective use of the lines; to the USDA-supported Maize Genetics Cooperation Stock Center, for distributing seed for *Ds-GFP* stocks; and to MaizeGDB for providing the informatic resources to identify relevant stocks. Our thanks also to federal employees at the US National Science Foundation, for their efforts to support, in as fair and transparent a manner as possible, the scientific community and its work to advance human understanding. We formatted this manuscript and the *SI Appendix* using and modifying a LATEX preprint template created by B. R. Keller (https://github.com/brenhinkeller/preprint-template.tex).

