## Supplementary material for "Successful Predictive Modeling of Pollen Fitness Phenotypes Is Enabled by Measures of Expression Specificity": ModelTrainingDatasets: README.rtf

Contents: 

DataS7_Genome_Model_Training_Dataset.csv
	A tabular dataset where each row is a B73 V5 gene model and each column is a genomic or expression feature used for training the Genome Model.
	For additional information on dataset curation see Methods Section A-D.1.

DataS7_Expression_Model_Training_Dataset.csv
	A tabular dataset where each row is a B73 V5 gene model and each column is a genomic or expression feature used for training the Expression Model.
	For additional information on dataset curation see Methods Section A-D.1.

DataS7_Combination_Model_Training_Dataset.csv
	A tabular dataset where each row is a B73 V5 gene model and each column is a genomic or expression feature used for training the Combination Model.
	For additional information on dataset curation see Methods Section A-D.1.

All gene ids correspond to the B73 V5 reference genome. 
Files created November 2025
Contact: 
