## Supplementary material for "Successful Predictive Modeling of Pollen Fitness Phenotypes Is Enabled by Measures of Expression Specificity": ProjectCode: CodeS1.ipynb

```
# %% [markdown]
# Imports

The following imports are necessary for the subsequent code. Project analyses were run using Python version 3.10.12

# %% [code]
import os
import sys
import math
import itertools
import argparse
import textwrap

import numpy as np
import pandas as pd

import matplotlib.pyplot as plt
import matplotlib.cm as cm
import matplotlib.lines as mlines
from matplotlib.ticker import MaxNLocator, FuncFormatter
import seaborn as sns
%matplotlib inline

from joblib import Parallel, delayed
from sklearn.linear_model import LogisticRegression, LinearRegression
from sklearn.ensemble import RandomForestClassifier
from sklearn.tree import DecisionTreeClassifier
from sklearn.model_selection import (
    train_test_split,
    StratifiedKFold,
    cross_val_score,
)
from sklearn import preprocessing
from sklearn.metrics import (
    accuracy_score,
    confusion_matrix,
    f1_score,
    recall_score,
    roc_auc_score,
    roc_curve,
    auc,
    r2_score,
)

from statsmodels.stats.multitest import multipletests

# plotting style 
sns.set_style("whitegrid")
plt.rcParams.update({
    "font.size": 11,
    "axes.titlesize": 13,
    "axes.labelsize": 11,
    "xtick.labelsize": 13,
    "ytick.labelsize": 12,
    "legend.fontsize": 10,
    "axes.spines.top": False,
    "axes.spines.right": False,
    "pdf.fonttype": 42, 
    "ps.fonttype": 42,  
    "svg.fonttype": "none"  
})


# %% [markdown]
## Load and filter validated phenotype dataset

This code reads in the insertion/transmission dataset and assigns class labels.

The following code cell also includes custom functions for calculating features, such as our specificity features (IPM, PSR, and NPS)


# %% [code]
df_path = 'DataS5_Full_Input_Dataset.csv' # INSERT PATH TO A DATAFRAME CSV FILE
read_in_df = pd.read_csv(df_path)

# make deep copy for downstream analysis
df = read_in_df.copy(deep=True)

# create another deep copy to retain "unused" instances for downstream evaluation 
validated_df_with_heldout_instances = read_in_df.copy(deep=True)

# set cutoff for binarization
cutoff_threshold = .45

# create an unused range df and remove that unused range from the current working df
unused_range_df = df[df['estimated_transmission_percentage'].between(0.45, 0.48, inclusive='both')]
df = df[~df['estimated_transmission_percentage'].between(0.45, 0.48, inclusive='both')]

# binarize the outcome and make a new column
df['Transmission_rate_class'] = df['estimated_transmission_percentage'].apply(lambda x: 0 if x>= cutoff_threshold else 1)

pre_nan_drop_validated_df = df.copy(deep=True)

# Arrays containing column names for RNA-Seq and proteomic profiling data, used for custom feature calculation functions. Update these with the named columns in your data if not using SI Appendix Data 5
proteomic_nonpollen_columns = ['Internode_6_7_x', 'Internode_7_8_x',
    'run_2_4_mm_from_tip_of_ear_primordium',
    'run_6_8_mm_from_tip_of_ear_primordium', 'embryos_20DAP', 'EMBRYOS',
    'endosperm_12DAP', 'endosperm_crown', 'mature_female_spikelets',
    'Germi0_0tin_Kernels_2_DAI', 'SYMMETRICAL_DIVISION_ZONE',
    'Leaf_Zone_2_Stomatal_x', 'GROWTH_ZONE', 'MATURE_LEAF_TISSUE_leaf_8',
    'pericarp_aleurone', 'PR', 'Cortex', 'EZ', 'MZ', 'SR', 'silks',
    'Vegetative_Meristem_and_surrounding_Tissue']

pollen_proteomic_nonpollen_columns = ['Mature_pollen', 'Max_dSNAF', 'avg_protein_abundance', 'protein_breadth']

rna_nonpollen_columns = ['Internode_6_7_y','Internode_7_8_y','Ear_Primordium_2_4_mm', 'Ear_Primordium_6_8_mm','Embryo_20_DAP','Embryo_38_DAP','Endosperm_12_DAP','Endosperm_Crown_27_DAP',
'Female_Spikelet_Collected_on_day_as_silk', 'Germinating_kernels_2_DAI','Leaf_Zone_1_Symmetrical','Leaf_Zone_2_Stomatal_y','Leaf_Zone_3_Growth','Mature_Leaf_8','Pericarp_Aleurone_27_DAP',
'Primary_Root_5_Days','Root_Cortex_5_Days','Root_Elongation_Zone_5_Days','Root_Meristem_Zone_5_Days','Secondary_Root_7_8_Days','Silk','Vegetative_Meristem_16_19_Days']

pollen_rna_columns = ['Bicellular_Male_Gametophyte', 'Microspore', 'Sperm_Cell', 'Anthers_R1', 'Mature_Pollen_paired_with_Sperm_Cells', 'Mature_Pollen_Walley', 'SpermCells_Chen', 'Max_FPKM']

pollen_protein_column = 'Mature_pollen'

pollen_rna_column = 'Mature_Pollen_Walley'

# see Walley et al. (2016) for a detailed explanation of derivation for 'max' features. This feature looks across all tissues expression values for a given gene id and is the max value
max_dNSAF = 'Max_dSNAF' 
max_FPKM = 'Max_FPKM'


# function for calculating non-pollen max protein expression. Update global arrays proteomic_column and pollen_proteomic_nonpollen_columns with your own proteomic expression tissue columns
def find_non_pollen_max_dsnaf(df_row):
    '''Function will find max dSNAF value that is not pollen by iterating df row argument columns and finding max'''

    non_pollen_max_dsnaf = None

    for col in df_row.index:
        if col in proteomic_nonpollen_columns and col not in pollen_proteomic_nonpollen_columns:
            
            if non_pollen_max_dsnaf is None:
                non_pollen_max_dsnaf = df_row[col]
            
            elif df_row[col] >= non_pollen_max_dsnaf:
                non_pollen_max_dsnaf = df_row[col]
        
    return non_pollen_max_dsnaf

# Function for calculating max non-pollen RNA-Seq. Update global array for rna_nonpollen_columns with your own RNA-Seq non-pollen tissue column names
def find_RNA_max_non_pollen_FPKM(df_row):
    '''Function will find max non-pollen rna-seq values by iterating df row argument columns and finding max'''
    
    non_pollen_max_fpkm = None
    rna_feature_list = rna_nonpollen_columns

    for col in df_row.index:
        if col in rna_feature_list:
            
            if non_pollen_max_fpkm is None:
                non_pollen_max_fpkm = df_row[col]
            
            elif df_row[col] >= non_pollen_max_fpkm:
                non_pollen_max_fpkm = df_row[col]
        
    return non_pollen_max_fpkm


# function for calculating IPM_P. Update global variable pollen_protein_column and max_dNSAF with your proteomic profiling column names
def is_max_in_pollen(df_row):
    '''Function finds if pollen is the maximally expressed tissue and returns 1 or 0 (T or F) for proteomic profiling'''
    try:
        if df_row[pollen_protein_column] == df_row[max_dNSAF]:
            return 1
        else:
            return 0
    except KeyError:
        if df_row[pollen_protein_column] == df_row[max_dNSAF]:
            return 1
        else:
            return 0

# function for calculating IPM_R. Update global variable rna_nonpollen_columns and pollen_rna_column with your own RNA-Seq column names
def is_max_FPKM_in_pollen(df_row):
        '''Function finds if pollen is the maximally expressed tissue and returns 1 or 0 (T or F) for RNA-Seq'''

        rna_feature_list = rna_nonpollen_columns + [pollen_rna_column]

        max_val = 0
        for col in df_row.index: 
            if col in rna_feature_list:
                if df_row[col] > max_val:
                    max_val = df_row[col]
        
        if max_val == df_row[pollen_rna_column]:
            return 1
        else:
            return 0

# function for calculating NPS_P. Update global variable proteomic_nonpollen_columns with your own proteomic profiling column names
def find_tissue_spec(df_row):
    """Function calculates the specificity ratio of the highest non-pollen tissue for proteomic profiling"""

    prot_feature_list = proteomic_nonpollen_columns

    temp_dict = {}
    for feat in prot_feature_list:
        
        row_max = 0
        for feat1 in prot_feature_list:
            
            if feat1 == feat:
                continue
            else:
                if df_row[feat1] > row_max:
                    row_max = df_row[feat1]

        ratio = (df_row[feat] + .001) / (row_max + .001)
        temp_dict[feat] = ratio

    return max(temp_dict.values())

# function to find NPS_R. Update global variable rna_nonpollen_columns and pollen_rna_column with your own RNA-Seq column names
def find_RNA_tissue_spec(df_row):
    """Function finds max non-pollen specificity ratio for RNA-Seq"""
    
    rna_feature_list = rna_nonpollen_columns

    temp_dict = {}
    for feat in rna_feature_list:
        
        row_max = 0
        for feat1 in rna_feature_list:
            
            if feat1 == feat:
                continue
            else:
                if df_row[feat1] > row_max:
                    row_max = df_row[feat1]

        ratio = (df_row[feat] + .001) / (row_max + .001)
        temp_dict[feat] = ratio

    return max(temp_dict.values())
print 

# calculate features
df['Max_Non-Pollen_dNSAF(protein)'] = df.apply(find_non_pollen_max_dsnaf, axis=1)
df['Max_Non-Pollen_FPKM'] = df.apply(find_RNA_max_non_pollen_FPKM, axis=1)
df['IPM_P'] = df.apply(is_max_in_pollen, axis=1)
df['IPM_R'] = df.apply(is_max_FPKM_in_pollen, axis=1)
df['PSR_P'] = (df[pollen_protein_column] + .1) / (df['Max_Non-Pollen_dNSAF(protein)']+ .1)
df['PSR_R'] = (df[pollen_rna_column] + .000001) / (df['Max_Non-Pollen_FPKM'] + .000001)
df['NPS_P'] = df.apply(find_tissue_spec, axis=1)
df['NPS_R'] = df.apply(find_RNA_tissue_spec, axis=1)
df['log(PSR_P)'] = df['PSR_P'].apply(lambda x: np.log(x))

# subset the full feature df for further ML modeling. sample columns shown here; can update with other feature columns 
# if using project datasets, make sure to include 'Gene_ID', 'feature_of_interest_ID', 'Transmission_rate_class'
df = df[['Gene_ID', 'feature_of_interest_ID', 'Transmission_rate_class',
    'Max_dSNAF', 'Max_FPKM', 'NPS_P']]
df.dropna(inplace=True)

# %% [markdown]
## Train model and evaluate performance

The following code prepares the features, labels, and CV partitions for classification modeling.

Assign custom weight values and regularization strength parameter C. Use custom weighting when a minority class is present. 

This code uses 3-fold stratified cross-validation to estimate auROC, sensitivity, and specificity.

Use this code to train the model and identify optimal class weight and C value. You can also use this code to compute Youden's index value. Alternatively, use the code in "CV Seed Test" cell to calculate avg. Youden's index. 


# %% [code]
# Project specific column drops and variable assignments. For X, make sure to remove any non-feature columns, y must be your phenotypic outcome variable. 
# Save identifying columns into variable id_columns for downstream analysis
X = df.drop(columns=['Transmission_rate_class', 'feature_of_interest_ID', 'Gene_ID'], axis=1)
y = df['Transmission_rate_class']
id_columns = df[['feature_of_interest_ID', 'Gene_ID']]

# normalize
X = (X-X.min())/(X.max()-X.min())
X.replace(np.nan, 0, inplace=True)

weights = {0:1, 1:10}

logregl1 = LogisticRegression(class_weight=weights, penalty='l1', solver='liblinear', C =5, random_state=42)
model = logregl1
kf = StratifiedKFold(n_splits=3, random_state=45, shuffle=True)

# metric variable assignments
best_fold = None
best_test_auc = 0
train_auc_scores = []
test_auc_scores = []
accuracy_scores = []
sensitivity_scores = []
specificity_scores = []
false_positive_rates = []
total_posi = []
pos_pred = []
total_nega = []
neg_pred = []
pos_class_df = pd.DataFrame()
neg_class_df = pd.DataFrame()
fpr_test_list = []
tpr_test_list =[]
best_test_fpr = None
best_test_tpr = None
youden_indicies = []
best_youden = None
plt.figure(figsize=(10, 8))

# Color maps for plots
colors_train = cm.Blues(np.linspace(0.4, 1, kf.n_splits))
colors_test = cm.Greens(np.linspace(0.4, 1, kf.n_splits))

# model training and testing
for fold, (train_index, test_index) in enumerate(kf.split(X, y), start=1):
    X_train, X_test = X.iloc[train_index], X.iloc[test_index]
    y_train, y_test = y.iloc[train_index], y.iloc[test_index]

    model.fit(X_train, y_train)

    y_train_prob = model.predict_proba(X_train)[:, 1]
    y_test_prob = model.predict_proba(X_test)[:, 1]

    train_auc = roc_auc_score(y_train, y_train_prob)
    test_auc = roc_auc_score(y_test, y_test_prob)

    train_auc_scores.append(train_auc)
    test_auc_scores.append(test_auc)
    
    fpr_train, tpr_train, thresholds_train = roc_curve(y_train, y_train_prob)
    fpr_test, tpr_test, thresholds_test = roc_curve(y_test, y_test_prob)

    fpr_test_list.append(fpr_test)
    tpr_test_list.append(tpr_test)

    youden_index = tpr_test - fpr_test
    optimal_idx = youden_index.argmax()
    optimal_threshold = thresholds_test[optimal_idx]
    
    plt.plot(fpr_train, tpr_train, color=colors_train[fold-1], lw=2, label=f'Fold {fold} Train ROC (AUC = {train_auc:.3f})')
    plt.plot(fpr_test, tpr_test, color=colors_test[fold-1], lw=2, label=f'Fold {fold} Test ROC (AUC = {test_auc:.3f})')

    y_test_pred = (y_test_prob >= optimal_threshold).astype(int)

    accuracy = accuracy_score(y_test, y_test_pred)
    sensitivity = recall_score(y_test, y_test_pred)
    tn, fp, fn, tp = confusion_matrix(y_test, y_test_pred).ravel()
    specificity = tn / (tn + fp)
    false_positive_rate = fp / (fp + tn)
    total_pos = tp + fn
    total_neg = tn + fp

    fold_pos_class_df = pd.DataFrame()
    fold_pos_class_df['Gene_ID'] = id_columns['Gene_ID'].iloc[test_index].values
    fold_pos_class_df['Transmission_rate_class'] = y.iloc[test_index].values
    fold_pos_class_df["predicted_class"] = y_test_pred

    fold_pos_class_df = fold_pos_class_df[fold_pos_class_df['Transmission_rate_class'] == 1]
    pos_class_df = pd.concat([pos_class_df, fold_pos_class_df])

    fold_neg_class_df = pd.DataFrame()
    fold_neg_class_df['Gene_ID'] = id_columns['Gene_ID'].iloc[test_index].values
    fold_neg_class_df['Transmission_rate_class'] = y.iloc[test_index].values
    fold_neg_class_df["predicted_class"] = y_test_pred

    fold_neg_class_df = fold_neg_class_df[fold_neg_class_df['Transmission_rate_class'] == 0]
    neg_class_df = pd.concat([neg_class_df, fold_neg_class_df])

    accuracy_scores.append(accuracy)
    sensitivity_scores.append(sensitivity)
    specificity_scores.append(specificity)
    false_positive_rates.append(false_positive_rate)
    total_posi.append(total_pos)
    pos_pred.append(tp)
    total_nega.append(total_neg)
    neg_pred.append(tn)
    youden_indicies.append(optimal_threshold)

    print(f"Fold {fold} - Train AUC = {train_auc:.3f}, Test AUC = {test_auc:.3f} \n")
    print(f"Test Fold {fold} - Accuracy = {accuracy:.3f}, Sensitivity = {sensitivity:.3f}, Specificity = {specificity:.3f}, False Positive Rate = {false_positive_rate:.3f} \n")
    print(f"Test Fold {fold} - correctly predicted {tp} positives out of {total_pos} positives\n")
    print("-----------------------------------------------------------------------------------\n")

    if test_auc > best_test_auc:
        best_test_auc = test_auc
        best_fold = fold
        best_model = model
        best_test_fpr = fpr_test
        best_test_tpr = tpr_test
        best_youden = optimal_threshold

plt.plot([0, 1], [0, 1], color='red', linestyle='--', lw=2, label='1:1 Line')

# ROC Plot
plt.xlim([0.0, 1.0])
plt.ylim([0.0, 1.05])
plt.xlabel('False Positive Rate', fontsize=20, fontweight = 'bold')
plt.ylabel('True Positive Rate', fontsize=20, fontweight = 'bold')
plt.title('ROC Curves for Train and Test by Fold')
plt.legend(loc='lower right', fontsize=16)
plt.grid(True)
plt.show()

print(f'Train_AUC_avg = {np.mean(train_auc_scores):.3f}, Test_AUC_avg = {np.mean(test_auc_scores):.3f}')
print(f'Accuracy_avg = {np.mean(accuracy_scores):.3f}, Sensitivity_avg = {np.mean(sensitivity_scores):.3f}, Specificity_avg = {np.mean(specificity_scores):.3f}, False_Positive_Rate_avg = {np.mean(false_positive_rates):.3f}')

avg_train_auc = np.mean(train_auc_scores)
avg_test_auc = np.mean(test_auc_scores)
avg_accuracy = np.mean(accuracy_scores)
avg_sensitivity = np.mean(sensitivity_scores)
avg_specificity = np.mean(specificity_scores)
avg_false_positive_rate = np.mean(false_positive_rates)

fig, ax = plt.subplots()
bar_width = 0.35
index = np.arange(len(train_auc_scores))

bar1 = ax.bar(index, train_auc_scores, bar_width, label='Train AUC')
bar2 = ax.bar(index + bar_width, test_auc_scores, bar_width, label='Test AUC')

ax.set_ylabel('AUC Value')
ax.set_title('Train and Test AUC Scores by CV Fold')
ax.set_xticks(index + bar_width / 2)
ax.set_xticklabels([f'Fold {i+1}' for i in range(len(train_auc_scores))])
ax.set_ylim(0, 1)
ax.set_yticks(np.arange(0, 1.1, 0.1))
ax.grid(axis='y', linestyle='--', linewidth=0.7)
ax.legend()

plt.tight_layout()
plt.show()

pd.set_option('display.max_rows', None)
np.set_printoptions(precision=15)
coef_weights = best_model.coef_[0]
features = X.columns
coef_df = pd.DataFrame({'Feature': features, 'Coefficient Weight': coef_weights})
coef_df = coef_df.sort_values(by='Coefficient Weight', ascending=False)
print("Coefficient Weights:")
coef_df


# %% [markdown]
## Genome-wide prediction for all model types

For each of the final trained models, load the corresponding training dataset and apply the chosen hyperparameters (C, class weight, Youden's index) identified in the previous cell.

The code will: 

  Refit the logistic regression model using the entire dataset as training data.
    
  Apply the model to predict outcomes on the genome-wide dataset, excluding the gene IDs used in the training set.
    
  Record, per gene, the:
   predicted probability of strong fitness effect,
   binary classification at the model's Youden's threshold (1 = predicted strong fitness effect, 0 = predicted no fitness effect)

  Merge predictions from all models into a single table.

The code also aligns all input features so that each gene in the genome-wide dataset is exported with its predictions and corresponding feature values. This combined table for the project models is provided as SI Appendix Data S9.

The data in DataS8_Genome_Wide_Dataset.csv is sourced from: 

S Sen, MR Woodhouse, JL Portwood, 2nd, CM Andorf, Maize feature store: A centralized resource to manage and analyze72
curated maize multi-omics features for machine learning applications. Database (Oxford) 2023 (2023)

MB Hufford, et al., De novo assembly, annotation, and comparative analysis of 26 diverse maize genomes. Science 373,84
655–662 (2021).


# %% [code]
# insert model name; update path to wrangled dataset. Use C, weight, and Youden's index chosen during model training
model_configs = [
    {   # for project specific Genome Model use C = 5, weight = 15, Youden's = .6976
        "name": "Genome Model",
        "csv_path": "DataS7_Model_Training_Datasets/DataS7_Genome_Model_Training_Datasets.csv",
        "C": 5,
        "weight": 15,
        "youden": .6976,
    },
    {   # for project specific Expression Model use C = 5, weight = 10, Youden's = .7609
        "name": "Expression Model",
        "csv_path": "DataS7_Model_Training_Datasets/DataS7_Expression_Model_Training_Datasets.csv",
        "C": 5,
        "weight": 10,
        "youden": 0.7609,
    },
    {   # for project specific Combination Model use C = 5, weight = 10, Youden's = .8118
        "name": "Combination Model",
        "csv_path": "DataS7_Model_Training_Datasets/DataS7_Combination_Model_Training_Dataset.csv",
        "C": 5,
        "weight": 10,
        "youden": 0.8118,
    }
]

# load full genome dataset
full_genome_path = 'DataS8_Genome_Wide_Dataset.csv' #insert path here
df_genome_full = pd.read_csv(full_genome_path)

# output variables/arrays 
merged_predictions = None
all_used_features = set()

# Compute specificity features
df_genome_full['log(PSR_P)'] = np.log(df_genome_full['PSR_P'])
df_genome_full['NPS_R'] = df_genome_full.apply(find_RNA_tissue_spec, axis=1)

# one hot encode categorical variables (project specific; update with any column names of new catagorical variables, or remove if your df does not have these categorical variables)
cat_features = ['subgenome_combine_final', 'Hufford2021_paired_status_final', "Subcel1_Wolf", "Localization_DeepLoc"]
df_genome_full = pd.get_dummies(df_genome_full, columns=cat_features)
df_genome_full = df_genome_full.applymap(lambda x: int(x) if isinstance(x, bool) else x)

for config in model_configs:
    print(f"\nprediction for: {config['name']}")

    # Load model-specific training data
    df_train = pd.read_csv(config["csv_path"])
    training_gene_ids = set(df_train["Gene_ID"])

    # Preprocess training data
    X_train = df_train.drop(columns=["Transmission_rate_class", "feature_of_interest_ID", "Gene_ID"])
    y_train = df_train["Transmission_rate_class"]
    X_train_copy = X_train.copy(deep=True)
    all_used_features.update(X_train_copy.columns)

    X_train = (X_train - X_train.min()) / (X_train.max() - X_train.min())
    X_train.replace(np.nan, 0, inplace=True)

    # Filter genome data to exclude the training set gene ids
    df_genome = df_genome_full[~df_genome_full['Gene_ID'].isin(training_gene_ids)].copy()

    # Project specific cleaning and renaming of columns for clarity
    X_genome = df_genome.drop(columns=['feature_of_interest_ID', 'Gene_ID'], errors='ignore')
    X_genome.rename(columns={
        'gene_breadth': 'gene_breadth(RNA_Seq)',
        'Anthers_R1': 'Anthers_R1(RNA_Seq)',
        'Mature_Pollen_Walley': 'Mature_Pollen_Walley(RNA_Seq)',
        'Max_dSNAF': 'Max_dNSAF(protein)',
        'Max_FPKM': 'Max_FPKM(RNA_Seq)'
    }, inplace=True)

    # Align features
    X_genome = X_genome[X_train_copy.columns]
    gene_ids = df_genome.loc[X_genome.index, 'Gene_ID'].values

    # normalize 
    X_genome = (X_genome - X_train_copy.min()) / (X_train_copy.max() - X_train_copy.min())
    X_genome.replace(np.nan, 0, inplace=True)

    # Train and predict
    model = LogisticRegression(
        class_weight={0: 1, 1: config["weight"]},
        penalty='l1',
        solver='liblinear',
        C=config["C"],
        random_state=42,
    )

    model.fit(X_train, y_train)
    
    probs_train = model.predict_proba(X_train)[:, 1]
    preds_train = (probs_train >= config["youden"]).astype(int)
    auroc = roc_auc_score(y_train, probs_train)
    cm = confusion_matrix(y_train, preds_train)
    tn, fp, fn, tp = cm.ravel()
    sensitivity = tp / (tp + fn) if (tp + fn) > 0 else np.nan
    specificity = tn / (tn + fp) if (tn + fp) > 0 else np.nan

    print(f"  Training Set AUROC: {auroc:.4f}")
    print(f"  Training Set Sensitivity (Recall for class 1): {sensitivity:.4f}")
    print(f"  Training Set Specificity (Recall for class 0): {specificity:.4f}")

    probs = model.predict_proba(X_genome)[:, 1]
    preds = (probs >= config["youden"]).astype(int)

    # predictions
    pred_df = pd.DataFrame({
        "Gene_ID": gene_ids,
        f"{config['name']}_Prob": probs,
        f"{config['name']}_Pred": preds,
    })

    # Merge
    if merged_predictions is None:
        merged_predictions = pred_df
    else:
        merged_predictions = pd.merge(merged_predictions, pred_df, on="Gene_ID", how="outer")


# Re-process full genome to extract all used features
df_genome_features = df_genome_full.copy()

# project specific column renaming 
df_genome_features.rename(columns={
    'gene_breadth': 'gene_breadth(RNA_Seq)',
    'Anthers_R1': 'Anthers_R1(RNA_Seq)',
    'Mature_Pollen_Walley': 'Mature_Pollen_Walley(RNA_Seq)',
    'Max_dSNAF': 'Max_dNSAF(protein)',
    'Max_FPKM': 'Max_FPKM(RNA_Seq)'
}, inplace=True)
df_genome_features['log(PSR_P)'] = np.log(df_genome_features['PSR_P'])
df_genome_features['NPS_R'] = df_genome_features.apply(find_RNA_tissue_spec, axis=1)
df_genome_features = df_genome_features.applymap(lambda x: int(x) if isinstance(x, bool) else x)

# Subset to contain features used across all models
common_features = list(all_used_features.intersection(df_genome_features.columns))
df_features_subset = df_genome_features[["Gene_ID"] + common_features]

# Final merge and save
final_df = pd.merge(merged_predictions, df_features_subset, on="Gene_ID", how="left")
final_df.to_csv("GenomeWideOutput.csv", index=False)

final_df
model_pred_cols = [col for col in final_df.columns if col.endswith('_Pred')]

# print summaries of predictions
for col in model_pred_cols:
    counts = final_df[col].value_counts(dropna=False).sort_index()
    print(f"\n{col}:")
    for val in [0, 1, np.nan]:
        if val in counts:
            label = "0" if val == 0 else "1" if val == 1 else "NaN"
            print(f"  {label}: {counts[val]}")

# %% [markdown]
## Load BonnMu insertion dataset and compute per-gene insertion counts

This code will read in the BonnMu insertion mutation dataset (DataS10_BonnMu_Dataset.csv) and perform our "BonnMu analysis."

Specifically, it calculates how many insertions were recovered in each genomic section.

The code will collapse the raw insertion calls into a per-gene count table (`partition_counts_df`) for each genomic section.

Subsequently, the BonnMu per-gene id insertion counts are merged with the genome-wide predictions. 

This allows the predicted average class difference to be computed for each genomic section using the full genome dataset.

The code also performs a permutation test to assess whether the observed mean insertion count difference for each genomic section is larger than expected by chance.

DataS10_BonnMu_Dataset.csv insertion line data is sourced from: 

C Marcon, et al., Bonnmu: A sequence-indexed resource of transposon-induced maize mutations for functional genomics84
studies . Plant Physiol. 184, 620–631 (2020)

YN Win, et al., Expanding the bonnmu sequence-indexed repository of transposon induced maize (zea mays l.) mutations86
in dent and flint germplasm. The Plant J. 120, 2253–2268 (2024).


# %% [code]
# Load the BonnMu dataset
BonnMu_dataset_path = 'DataS10_BonnMu_Dataset.csv' # insert path here
BonnMu_df = pd.read_csv(BonnMu_dataset_path, delimiter='\t')
BonnMu_df = BonnMu_df[['Gene_ID', 'partition', 'InsertionIdentifier']].drop_duplicates()

# Count partitions per gene
partition_counts_df = BonnMu_df.groupby(['Gene_ID'])['partition'].value_counts().unstack(fill_value=0).reset_index()

desired_partition_order = [
    "promoterProx",
    "promoterCore",
    "five_prime_utr",
    "exon",
    "intron",
    "three_prime_utr",
    "putative close downstream enhancer",
]

desired_partition_order = [i for i in desired_partition_order if i in partition_counts_df.columns]

partition_labels = {
    "promoterProx": "Promoter \n Proximal",
    "promoterCore": "Promoter Core",
    "five_prime_utr": "5\u2032 UTR",    
    "three_prime_utr": "3\u2032 UTR",     
    "exon": "Exon",
    "intron": "Intron",
    "putative close downstream enhancer": "Downstream",
}

model_titles = {
    "Genome_Model_2_Pred": "Genome Model",
    "Expression_Model_Pred": "Expression Model",
    "Combination_Model_Pred": "Combination Model"
}


def model_title_clean(col):
    name = col.replace("_Pred", "")
    name = name.replace("_", " ")
    # Tidy common tokens:
    name = name.replace("auroc", "auROC").replace("utr", "UTR")
    return name.title()


def partition_clean(key):
    return wrap_label(partition_labels.get(key, key.replace("_", " ")))

def wrap_label(s, width=18):
    return "\n".join(textwrap.wrap(s, width=width))

# helper function for p-value printing 
def p_to_stars(p):
    if p < 1e-4: return "****"
    if p < 1e-3: return "***"
    if p < 1e-2: return "**"
    if p < 5e-2: return "*"
    return "n.s."

# helper function to draw a bracket for the p-value on figure
def add_sig_bracket(ax, x1, x2, y, h, text, lw=1.2):
    """Draw a significance bracket between x1 and x2 at height y with bump h."""
    ax.plot([x1, x1, x2, x2], [y, y+h, y+h, y], linewidth=lw, color='black')
    ax.text((x1+x2)/2, y+h*1.1, text, ha='center', va='bottom', fontsize=13, color='black')

# BonnMu analysis
def compute_partition_diff(pred_labels, gene_ids, partition_counts_df, desired_partition_order):
    shuffled_df = pd.DataFrame({
        'Gene_ID': gene_ids,
        'Shuffled_Pred': pred_labels
    })

    pos_ids = shuffled_df[shuffled_df['Shuffled_Pred'] == 1]['Gene_ID']
    neg_ids = shuffled_df[shuffled_df['Shuffled_Pred'] == 0]['Gene_ID']

    pos_subset = pd.merge(pos_ids, partition_counts_df, on='Gene_ID', how='left').dropna()
    neg_subset = pd.merge(neg_ids, partition_counts_df, on='Gene_ID', how='left').dropna()

    if len(pos_subset) == 0 or len(neg_subset) == 0:
        return np.zeros(len(desired_partition_order))

    # calculate the means 
    pos_norm = pos_subset[desired_partition_order].sum() / len(pos_subset)
    neg_norm = neg_subset[desired_partition_order].sum() / len(neg_subset)

    return (pos_norm - neg_norm).values

# Run analysis for each prediction column
model_pred_cols = [col for col in final_df.columns if col.endswith('_Pred')]
combined_model_data = {}

for pred_col in model_pred_cols:
    pos_predictions = final_df[final_df[pred_col] == 1]
    neg_predictions = final_df[final_df[pred_col] == 0]

    pos_ids = pos_predictions['Gene_ID']
    neg_ids = neg_predictions['Gene_ID']

    BonnMu_pos_subset = pd.merge(pos_ids, partition_counts_df, on='Gene_ID', how='left').dropna()
    BonnMu_neg_subset = pd.merge(neg_ids, partition_counts_df, on='Gene_ID', how='left').dropna()

    BonnMu_pos_length = len(BonnMu_pos_subset)
    BonnMu_neg_length = len(BonnMu_neg_subset)

    BonnMu_pos_subset = BonnMu_pos_subset[desired_partition_order].agg('sum') / BonnMu_pos_length
    BonnMu_neg_subset = BonnMu_neg_subset[desired_partition_order].agg('sum') / BonnMu_neg_length

    combined_df = pd.DataFrame({
        'Positive': BonnMu_pos_subset,
        'Negative': BonnMu_neg_subset
    })

    combined_model_data[pred_col] = combined_df

    # set up permutation test
    np.random.seed(42) 
    observed_diff = combined_df['Positive'] - combined_df['Negative']  # per partition

    n_permutations = 1000
    permutation_diffs = np.zeros((n_permutations, len(desired_partition_order)))
    gene_ids = final_df['Gene_ID'].values
    true_preds = final_df[pred_col].values

    # Run permutation test
    for i in range(n_permutations):
        shuffled_preds = np.random.permutation(true_preds)
        permutation_diffs[i] = compute_partition_diff(
            shuffled_preds, gene_ids, partition_counts_df, desired_partition_order
        )

    pvals_one = []
    for j, part in enumerate(desired_partition_order):
        null_dist = permutation_diffs[:, j]
        obs = observed_diff[part]

        # One-tailed 
        p_one = np.mean(null_dist <= obs)

        # Avoid exactly zero p-value
        if p_one == 0:
            p_one = 1.0 / (n_permutations + 1)

        pvals_one.append(p_one)

    pval_one_series = pd.Series(pvals_one, index=desired_partition_order)
    print(f"\n ONE-tailed permutation p-values (Positive < Negative) for {pred_col}:\n")
    print(pval_one_series)

    # plot
    fig, ax = plt.subplots(figsize=(22/2.54, 18/2.54))
    categories = combined_df.index
    x = np.arange(len(categories))
    bar_width = 0.35

    neg_bars = ax.bar(x - bar_width/2, combined_df['Negative'], width=bar_width,
                    color='#6baed6', alpha=0.8, label='Predicted no fitness effect')
    pos_bars = ax.bar(x + bar_width/2, combined_df['Positive'], width=bar_width,
                    color='#fb6a4a', alpha=0.9, label=' Predicted strong fitness effect')
    ax.set_ylim(0,8.5)
   
    # plot insertion count bars 
    ax.set_xticks(list(x))
    pretty_xticks = [partition_clean(p) for p in categories]
    ax.set_xticklabels(pretty_xticks, rotation=45, ha='center')
    ax.set_ylabel("Mean insertions per genomic section", fontsize = 16)
    title = model_titles.get(pred_col, model_title_clean(pred_col))
    ax.set_title(f'{title}', pad=10, fontsize= 22)
    ax.yaxis.set_major_locator(MaxNLocator(nbins=5))
    ax.yaxis.set_major_formatter(FuncFormatter(lambda v, pos: f"{v:.1f}"))
    ax.grid(axis='y', linewidth=0.6, alpha=0.35)
    sns.despine(ax=ax)

    ax.legend(
    frameon=True,        
    facecolor="white",    
    framealpha=1.0,      
    fontsize=16,
    ncol=1,
    loc="upper right"
)
    #plot p-values
    ylim_top = ax.get_ylim()[1]
    ax.set_ylim(top=ylim_top * 1.15)
    for i, part in enumerate(desired_partition_order):
        y_max = max(neg_bars[i].get_height(), pos_bars[i].get_height())
        y = y_max * 1.02
        h = y_max * 0.035 if y_max > 0 else 0.035  
        x1 = i - bar_width/2
        x2 = i + bar_width/2
        p_adj = pval_one_series[part]
        stars = p_to_stars(p_adj)
        if p_adj == 0:
            p_text = "p<1e-3" if n_permutations >= 1000 else f"p<{1/n_permutations:.5f}"
        else:
            p_text = f"p={p_adj:.2g}"
            
        label = f"{stars}{p_text}" if stars != "n.s." else f"{p_text}"
        add_sig_bracket(ax, x1, x2, y, h, label)

    plt.tight_layout()
    # fig.savefig(f"{title} BonnMu Test.pdf", bbox_inches="tight", dpi=600)  
    plt.show()


# %% [markdown]
### Training set BonnmMu analysis

The following code performs the above "BonnMu analysis" on the training dataset. 

# %% [code]
# set up BonnMu analysis
gene_ids  = pre_nan_drop_validated_df['Gene_ID'].values
true_preds = pre_nan_drop_validated_df['Transmission_rate_class'].values

pos_predictions = pre_nan_drop_validated_df[pre_nan_drop_validated_df['Transmission_rate_class'] == 1]
neg_predictions = pre_nan_drop_validated_df[pre_nan_drop_validated_df['Transmission_rate_class'] == 0]

pos_ids = pos_predictions['Gene_ID']
neg_ids = neg_predictions['Gene_ID']

BonnMu_pos_subset = pd.merge(pos_ids, partition_counts_df, on='Gene_ID', how='left').dropna()
BonnMu_neg_subset = pd.merge(neg_ids, partition_counts_df, on='Gene_ID', how='left').dropna()

BonnMu_pos_length = len(BonnMu_pos_subset)
BonnMu_neg_length = len(BonnMu_neg_subset)

BonnMu_pos_subset = BonnMu_pos_subset[desired_partition_order].agg('sum') / max(BonnMu_pos_length, 1)
BonnMu_neg_subset = BonnMu_neg_subset[desired_partition_order].agg('sum') / max(BonnMu_neg_length, 1)

combined_df = pd.DataFrame({
    'Positive': BonnMu_pos_subset,
    'Negative': BonnMu_neg_subset
})

# set up permutation test
np.random.seed(42)
n_permutations = 1000
observed_diff = combined_df['Positive'] - combined_df['Negative']
permutation_diffs = np.zeros((n_permutations, len(desired_partition_order)))

for i in range(n_permutations):
    shuffled_preds = np.random.permutation(true_preds)
    permutation_diffs[i] = compute_partition_diff(
        shuffled_preds, gene_ids, partition_counts_df, desired_partition_order
    )

# one-tailed p-values
pvals_one = []
for j, part in enumerate(desired_partition_order):
    null_dist = permutation_diffs[:, j]
    obs = observed_diff[part]
    p_one = np.mean(null_dist <= obs) 
    if p_one == 0:
        p_one = 1.0 / (n_permutations + 1)  # avoid zero division
    pvals_one.append(p_one)

pval_one_series = pd.Series(pvals_one, index=desired_partition_order)

print("\nONE-tailed permutation p-values (Positive < Negative):\n")
print(pval_one_series.round(4))

# helper function for p-value printing 
def p_to_stars(p):
    if p < 1e-4: return "****"
    if p < 1e-3: return "***"
    if p < 1e-2: return "**"
    if p < 5e-2: return "*"
    return "n.s."

def add_sig_bracket(ax, x1, x2, y, h, text, lw=1.2):
    ax.plot([x1, x1, x2, x2], [y, y+h, y+h, y], linewidth=lw, color='black')
    ax.text((x1+x2)/2, y+h*1.1, text, ha='center', va='bottom', fontsize=13, color='black')


# plotting 
fig, ax = plt.subplots(figsize=(22/2.54, 18/2.54))
x = np.arange(len(desired_partition_order))
bar_width = 0.35

neg_bars = ax.bar(x - bar_width/2, combined_df['Negative'], width=bar_width,
                  color='#6baed6', alpha=0.8, label='No fitness effect')
pos_bars = ax.bar(x + bar_width/2, combined_df['Positive'], width=bar_width,
                  color='#fb6a4a', alpha=0.9, label='Strong fitness effect')

ax.set_ylim(0,8.5)
ax.set_xticks(list(x))
ax.set_xticklabels(pretty_xticks, rotation=45, ha='center')
ax.set_ylabel("Mean insertions per genomic section", fontsize = 16)
ax.set_title("Training Dataset", pad=10, fontsize = 22)
ax.legend(
    frameon=True,       
    facecolor="white",   
    framealpha=1.0,     
    fontsize=16,
    ncol=1,
    loc="upper right"
)
ax.yaxis.set_major_locator(MaxNLocator(nbins=5))
ax.yaxis.set_major_formatter(FuncFormatter(lambda v, pos: f"{v:.1f}"))
ax.grid(axis='y', linewidth=0.6, alpha=0.35)
sns.despine(ax=ax)
ylim_top = ax.get_ylim()[1]
ax.set_ylim(top=ylim_top * 1.15)

for i, part in enumerate(desired_partition_order):
    y_max = max(neg_bars[i].get_height(), pos_bars[i].get_height())
    y = y_max * 1.02
    h = y_max * 0.035 if y_max > 0 else 0.035

    x1 = i - bar_width/2
    x2 = i + bar_width/2

    p_raw = pval_one_series[part]
    stars = p_to_stars(p_raw)
    p_text = "p<1e-3" if (p_raw == 0 and n_permutations >= 1000) else f"p={p_raw:.2g}"
    label = f"{stars}{p_text}" if stars != "n.s." else f"{p_text}"

    add_sig_bracket(ax, x1, x2, y, h, label)

plt.tight_layout()
# fig.savefig(f"Training Set_BonnMu Test.pdf", bbox_inches="tight", dpi=600)   
plt.show()


# %% [markdown]
## BonnMu analysis of predicted strong fitness effect

This code uses the genome-wide predicted strong fitness effect genes, split into an upper 50% and lower 50% based on predicted probability. 

The following analysis follows the same procedure as the "BonnMu analysis" performed above, except that instead of comparing mean insertion count differences between strong fitness effect and no fitness effect predictions, we take only genes predicted as strong fitness effect and split them into upper and lower 50% groups based on their predicted probabilities. The mean insertion count difference between the two groups is then evaluated. 


# %% [code]
# set up naming dictionaries and helper functions
partition_labels = {
    "promoterProx": "Promoter \n Proximal",
    "promoterCore": "Promoter Core",
    "five_prime_utr": "5′ UTR",
    "three_prime_utr": "3′ UTR",
    "exon": "Exon",
    "intron": "Intron",
    "putative close downstream enhancer": "Downstream"
}
model_titles = {
    "Genome_Model_2_Pred": "Genome Model",
    "Expression_Model_Pred": "Expression Model",
    "Boost_Model_Pred": "Combination Model"
}

def model_title_clean(col):
    name = col.replace("_Pred", "")
    name = name.replace("_", " ")
    name = name.replace("auroc", "auROC").replace("utr", "UTR")
    return name.title()

col_lower = "#9ecae1"  
col_upper = "#3182bd"  

# helper function for p-value printing 
def p_to_stars(p):
    if p < 1e-4: return "****"
    if p < 1e-3: return "***"
    if p < 1e-2: return "**"
    if p < 5e-2: return "*"
    return "n.s."

def add_sig_bracket(ax, x1, x2, y, h, text, lw=1.2):
    """Draw a significance bracket between x1 and x2 at height y with bump h."""
    ax.plot([x1, x1, x2, x2], [y, y+h, y+h, y], linewidth=lw, color='black')
    ax.text((x1+x2)/2, y+h*1.1, text, ha='center', va='bottom', fontsize=13, color='black')

def norm_partitions(df_ids, partition_counts_df, desired_partition_order):
    """Mean insertions per gene for a set of Gene_IDs. This returns a series indexed by desired_partition_order."""
    if df_ids.empty:
        return pd.Series(0.0, index=desired_partition_order)
    merged = (
        pd.merge(df_ids['Gene_ID'], partition_counts_df, on='Gene_ID', how='left')
        .dropna()
    )
    if merged.empty:
        return pd.Series(0.0, index=desired_partition_order)
    return merged[desired_partition_order].sum() / len(merged)

def permuted_upper_minus_lower(pos_gene_ids, pos_probs, midpoint,
                               partition_counts_df, desired_partition_order, rng):
    """Shuffle probs among positives, split by prob rank, return (Upper - Lower) for partitions."""

    order = rng.permutation(len(pos_probs))
    shuffled_probs = pos_probs[order]

    
    sort_idx = np.argsort(shuffled_probs)
    lower_idx = sort_idx[:midpoint]
    upper_idx = sort_idx[midpoint:]

    lower_ids = pd.DataFrame({'Gene_ID': pos_gene_ids[lower_idx]})
    upper_ids = pd.DataFrame({'Gene_ID': pos_gene_ids[upper_idx]})

    lower_part = norm_partitions(lower_ids, partition_counts_df, desired_partition_order)
    upper_part = norm_partitions(upper_ids, partition_counts_df, desired_partition_order)

    return (upper_part - lower_part).values


model_pred_cols = [col for col in final_df.columns if col.endswith('_Pred')]
model_prob_cols = [col.replace('_Pred', '_Prob') for col in model_pred_cols]

for pred_col, prob_col in zip(model_pred_cols, model_prob_cols):
  
    if prob_col not in final_df.columns:
        print(f"Missing probability column for {pred_col}: {prob_col}")
        continue

    print(f"Processing model: {pred_col}")

    pos_predictions = (
        final_df.loc[final_df[pred_col] == 1, ['Gene_ID', prob_col]]
        .dropna(subset=[prob_col])
        .copy()
    )

    if pos_predictions.empty:
        print(f"No positive predictions for {pred_col}")
        continue

    # Split into lower and upper halves
    sorted_pos = pos_predictions.sort_values(by=prob_col, kind="mergesort") 
    midpoint = len(sorted_pos) // 2
    lower_half = sorted_pos.iloc[:midpoint]
    upper_half = sorted_pos.iloc[midpoint:]
    print(len(lower_half), len(upper_half))


    # Merge with BonnMu partitions and normalize
    lower_partitions = norm_partitions(lower_half, partition_counts_df, desired_partition_order)
    upper_partitions = norm_partitions(upper_half, partition_counts_df, desired_partition_order)

    # Build combined df
    combined_df = pd.DataFrame({
        'Lower 50%': lower_partitions,
        'Upper 50%': upper_partitions
    }).reindex(desired_partition_order)

    # clean x tick labels
    categories = combined_df.index
    pretty_xticks = [partition_clean(p) for p in categories]

    # difference calculation
    observed_diff = (combined_df['Upper 50%'] - combined_df['Lower 50%']).values

    n_permutations = 1000  # adjust if you want more precision
    rng = np.random.default_rng(seed=42)

    pos_gene_ids = pos_predictions['Gene_ID'].to_numpy()
    pos_probs = pos_predictions[prob_col].to_numpy()

    perm_diffs = np.zeros((n_permutations, len(desired_partition_order)))
    for i in range(n_permutations):
        perm_diffs[i, :] = permuted_upper_minus_lower(
            pos_gene_ids, pos_probs, midpoint,
            partition_counts_df, desired_partition_order, rng
        )

    # one-tailed p-values
    pvals_one = []
    for j in range(len(desired_partition_order)):
        null_j = perm_diffs[:, j]
        obs_j = observed_diff[j]
        p_one = np.mean(null_j <= obs_j)
        if p_one == 0:
            p_one = 1.0 / (n_permutations + 1) 
        pvals_one.append(p_one)
    pval_one_series = pd.Series(pvals_one, index=desired_partition_order)
    print(f"\nONE-tailed permutation p-values (Upper < Lower) for {pred_col}:")
    print(pval_one_series.round(4))

    # plotting
    fig, ax = plt.subplots(figsize=(22/2.54, 18/2.54))
    x = np.arange(len(combined_df))
    bar_width = 0.38

    bars_lower = ax.bar(x - bar_width/2, combined_df['Lower 50%'],
                        width=bar_width, color=col_lower, label='Lower 50%')
    bars_upper = ax.bar(x + bar_width/2, combined_df['Upper 50%'],
                        width=bar_width, color=col_upper, label='Upper 50%')

    title = model_titles.get(pred_col, model_title_clean(pred_col))
    ax.set_title(f"{title}", pad=10, fontsize = 22)
    ax.set_ylabel("Mean insertions per genomic section", fontsize = 16)
    ax.set_xticks(list(x))
    ax.set_xticklabels(pretty_xticks, rotation=45, ha='center')
    ax.yaxis.set_major_locator(MaxNLocator(nbins=5))
    ax.yaxis.set_major_formatter(FuncFormatter(lambda v, pos: f"{v:.1f}"))
    ax.grid(axis='y', linewidth=0.6, alpha=0.35)
    sns.despine(ax=ax)
    ax.legend(
    frameon=True,      
    facecolor="white",   
    framealpha=1.0,     
    fontsize=15,
    ncol=1,
    loc="upper right"
)
    ylim_lo, ylim_hi = ax.get_ylim()
    ax.set_ylim(ylim_lo, ylim_hi * 1.15)

    for i, part in enumerate(desired_partition_order):
        y_max = max(bars_lower[i].get_height(), bars_upper[i].get_height())
        y = y_max * 1.02
        h = max(0.03 * (ylim_hi if ylim_hi > 0 else 1), 0.02)  

        x1 = i - bar_width/2
        x2 = i + bar_width/2

        p_adj = pval_one_series[part]
        stars = p_to_stars(p_adj)
        p_text = f"p={p_adj:.2g}" if p_adj > 0 else f"p<{1/(n_permutations+1):.3f}"
        label = f"{stars}{p_text}" if stars != "n.s." else p_text
        add_sig_bracket(ax, x1, x2, y, h, label)

    fig.tight_layout()
    # fig.savefig(f"{title} BonnMu Positives Test.pdf", bbox_inches="tight", dpi=600)    
    plt.show

# %% [markdown]
### Feature Subset Test

The following code takes a model ready dataset (i.e., cleaned feature columns).

The code tests every combination of the provided features. You can adjust min and max combination sizes

Specifically, it runs each feature subset model across multiple CV seeds to estimate average performance at different C and class weight values.

It returns a CSV containing all tested feature subset combinations along with their average performance metrics.

See the inline comments for run specifics, such as your own dataset path and minimum and maximum subset sizes.

# %% [code]
csv_path = 'DataS7_Model_Training_Datasets/DataS7_Genome_Model_Training_Datasets.csv' # update with your dataset path. This should be a cleaned and wrangled dataset ready for modeling with feature columns of interest
C_values = [0.4, 1, 5] # specified C values to test
weight_values = [15] # specified weight values to test
n_jobs = 1 # parallel workers to use locally


df = pd.read_csv(csv_path)
X_full = df.drop(columns=["Transmission_rate_class", "feature_of_interest_ID", "Gene_ID"])
y = df["Transmission_rate_class"]

# normalize 
X_full = (X_full - X_full.min()) / (X_full.max() - X_full.min())
X_full.replace(np.nan, 0, inplace=True)
feature_list = list(X_full.columns)

# model evaluation function
def evaluate_feature_combo(selected_features, weight, C_value, seeds=150):
    '''This function evaluates a feature combination using stratified K-Fold cross-validation across specified random seeds.'''
    X = X_full[list(selected_features)]
    test_aurocs = []
    weights = {0: 1, 1: weight}

    for seed in range(seeds):
        kf = StratifiedKFold(n_splits=3, random_state=seed, shuffle=True)
        fold_aurocs = []

        for train_idx, test_idx in kf.split(X, y):
            X_train, X_test = X.iloc[train_idx], X.iloc[test_idx]
            y_train, y_test = y.iloc[train_idx], y.iloc[test_idx]

            model = LogisticRegression(
                class_weight=weights,
                penalty="l1",
                solver="liblinear",
                C=C_value,
                random_state=42,
            )
            model.fit(X_train, y_train)
            y_test_prob = model.predict_proba(X_test)[:, 1]
            fold_aurocs.append(roc_auc_score(y_test, y_test_prob))

        test_aurocs.append(np.mean(fold_aurocs))

    return np.mean(test_aurocs), np.std(test_aurocs)


def run_single_combo(feature_combo, weight, C_val):
    avg_auroc, std_auroc = evaluate_feature_combo(feature_combo, weight, C_val, seeds=150)
    return {
        "Avg_Seed_Test_auROC": avg_auroc,
        "Std_Seed_Test_auROC": std_auroc,
        "Features_Used": list(feature_combo),
        "Weight_and_C": f"Weight={weight}, C={C_val}",
    }


def run_with_counter(index, total, combo, weight, C_val):
    print(f"Running job {index+1}/{total}: features={combo}, weight={weight}, C={C_val}")
    result = run_single_combo(combo, weight, C_val)
    result["Job_Index"] = index + 1
    return result

# set up jobs and return information 
all_jobs_stub = [
    (feature_combo, weight, C_val)
    for k in range(1, 4) # custom range for min and max features in subsets (min must be at least 1)
    for feature_combo in itertools.combinations(feature_list, k)
    for weight in weight_values
    for C_val in C_values
]

total_jobs = len(all_jobs_stub)
all_jobs = [
    (i, total_jobs, combo, weight, C_val)
    for i, (combo, weight, C_val) in enumerate(all_jobs_stub)
]

print(f"Starting job loop with {total_jobs} total combos using {n_jobs} parallel workers")

# parallel execution
results = Parallel(n_jobs=n_jobs)(
    delayed(run_with_counter)(*job) for job in all_jobs
)

# print and save results
results_df = pd.DataFrame(results)
results_df = results_df.sort_values(by="Avg_Seed_Test_auROC", ascending=False)

output_filename = f"Feature_Subset_auROC_Results.csv"
results_df.to_csv(output_filename, index=False)
print(f"Saved results to: {output_filename}")

# %% [markdown]
### CV Seed Test

The following code performs a cross-validation seed test.

This code uses model ready datasets as defined in the model config dictionary in the genome-wide analysis code above. 

The code trains and evaluates the models across the specified seed range and returns the average performance metrics.

# %% [code]
all_results = []

# main run loop
for config in model_configs:
    print(f"Running {config['name']}")

    df = pd.read_csv(config["csv_path"])
    X = df.drop(
        columns=["Transmission_rate_class", "feature_of_interest_ID", "Gene_ID"]
    )
    y = df["Transmission_rate_class"]

    # Normalize and clean
    X = (X - X.min()) / (X.max() - X.min())
    X.replace(np.nan, 0, inplace=True)

    weights = {0: 1, 1: config["weight"]}
    model_C = config["C"]

    # function for seed evaluation
    def evaluate_seed(seed):
        kf = StratifiedKFold(n_splits=3, random_state=seed, shuffle=True)
        test_aurocs = []
        accs, senss, specs = [], [], []

        for train_idx, test_idx in kf.split(X, y):
            X_train, X_test = X.iloc[train_idx], X.iloc[test_idx]
            y_train, y_test = y.iloc[train_idx], y.iloc[test_idx]

            model = LogisticRegression(
                class_weight=weights,
                penalty="l1",
                solver="liblinear",
                C=model_C,
                random_state=42,
            )
            model.fit(X_train, y_train)

            y_test_prob = model.predict_proba(X_test)[:, 1]
            test_auc = roc_auc_score(y_test, y_test_prob)

            fpr, tpr, thresholds = roc_curve(y_test, y_test_prob)
            youden_index = np.argmax(tpr - fpr)
            optimal_threshold = thresholds[youden_index]

            y_pred = (y_test_prob >= optimal_threshold).astype(int)
            tn, fp, fn, tp = confusion_matrix(y_test, y_pred).ravel()

            acc = (tp + tn) / (tp + tn + fp + fn)
            sens = tp / (tp + fn) if (tp + fn) > 0 else 0   
            spec = tn / (tn + fp) if (tn + fp) > 0 else 0   

            test_aurocs.append(test_auc)
            accs.append(acc)
            senss.append(sens)
            specs.append(spec)

        return (
            np.mean(test_aurocs),
            np.mean(accs),
            np.mean(senss),
            np.mean(specs),
        )

    seeds = range(1, 1000)
    for seed in seeds:
        avg_auroc, avg_acc, avg_sens, avg_spec = evaluate_seed(seed)
        all_results.append(
            {
                "Seed": seed,
                "Avg_auROC": avg_auroc,
                "Avg_Accuracy": avg_acc,
                "Avg_Sensitivity": avg_sens,
                "Avg_Specificity": avg_spec,
                "Model": config["name"],
            }
        )

# final results saving and printing
results_df = pd.DataFrame(all_results)
results_df.to_csv("Seed_test_results.csv", index=False)

print("\n=== AVERAGE METRICS (MEAN ± STD) ACROSS 1000 SEEDS ===\n")
grouped_mean = results_df.groupby("Model").mean(numeric_only=True)
grouped_std = results_df.groupby("Model").std(numeric_only=True)

for model in grouped_mean.index:
    print(f"Model: {model}")
    print(f"  Avg auROC:       {grouped_mean.loc[model, 'Avg_auROC']:.4f} ± {grouped_std.loc[model, 'Avg_auROC']:.4f}")
    print(f"  Avg Accuracy:    {grouped_mean.loc[model, 'Avg_Accuracy']:.4f} ± {grouped_std.loc[model, 'Avg_Accuracy']:.4f}")
    print(f"  Avg Sensitivity: {grouped_mean.loc[model, 'Avg_Sensitivity']:.4f} ± {grouped_std.loc[model, 'Avg_Sensitivity']:.4f}")
    print(f"  Avg Specificity: {grouped_mean.loc[model, 'Avg_Specificity']:.4f} ± {grouped_std.loc[model, 'Avg_Specificity']:.4f}")
    print("-" * 40)

# plotting
plt.figure(figsize=(8.7,7.1))
ax = sns.boxplot(data=results_df, x="Model", y="Avg_auROC",
                 linewidth=2.0, color='skyblue')

# Compute means
x_positions = ax.get_xticks()
x_labels = results_df["Model"].unique()
means = results_df.groupby("Model")["Avg_auROC"].mean()

# yellow diamond mean markers and text labels
for idx, model_name in enumerate(x_labels):
    mean = means[model_name]
    ax.scatter(
        x_positions[idx], mean,
        color="yellow", edgecolor="black", linewidth=1.2,
        marker="D", s=50, zorder=10,
        label="Mean auROC" if idx == 0 else None
    )
    ax.text(
        x_positions[idx] + 0.05, mean - 0.002,
        f"{mean:.3f}", ha="left", va="center",
        fontsize=8, fontweight="bold", color="black", zorder=11
    )

ax.set_ylabel("Avg auROC", fontsize=12, fontweight="bold", labelpad=10)
ax.set_xlabel("", fontsize=12, fontweight="bold")

ax.set_xticklabels(ax.get_xticklabels(), rotation=45,
                   fontsize=10, fontweight="bold", ha="right")
ax.tick_params(axis='y', labelsize=10)

handles, labels = ax.get_legend_handles_labels()
by_label = dict(zip(labels, handles))
ax.legend(by_label.values(), by_label.keys(),
          fontsize=10, loc="upper center",
          frameon=True, edgecolor="black")

ax.grid(True, linestyle='--', linewidth=0.5, alpha=0.7)

plt.tight_layout()
plt.show()
```
