## Supporting_Information_and_Supplemental_File_Index for "Successful Predictive Modeling of Pollen Fitness Phenotypes Is Enabled by Measures of Expression Specificity"

### Supporting Information for: Successful Predictive Modeling of Pollen Fitness Phenotypes Is Enabled by Measures of Expression Specificity

Preprint, compiled January 13, 2026

**Sebastian A.F. Mueller, Zuzana Vejlupkova, Molly Megraw, John E. Fowler**

#### Supporting Information Text

#### 1 Overview of Machine Learning Modeling Process

##### ML Model Form

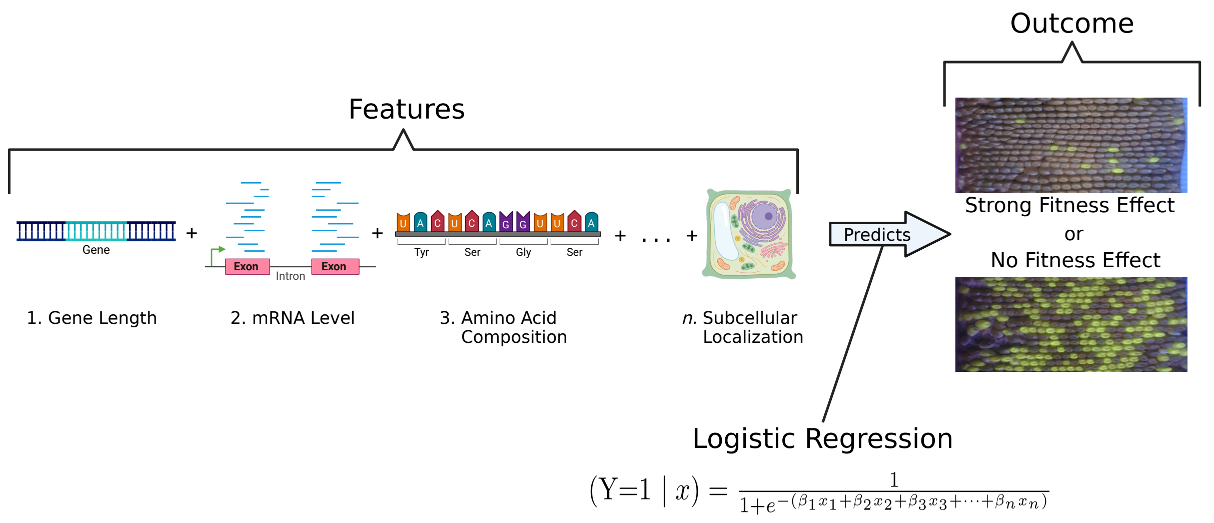

**Figure S1:** Machine Learning can integrate disparate data types to predict biological outcomes. In this study, models used information about single gene models, such as gene length or RNA-seq read count, to predict whether a mutation in the given gene is likely to cause a strong fitness effect or no fitness effect in pollen. Specifically, we applied logistic regression, which learns relationships between these features and the probability of a fitness effect outcome. Figure created in BioRender. Mueller, S. (2025) https://BioRender.com/vxwu4q2

**RNA-Seq vs Protein Expression Feature Predictive Strength Test**

**(A)**

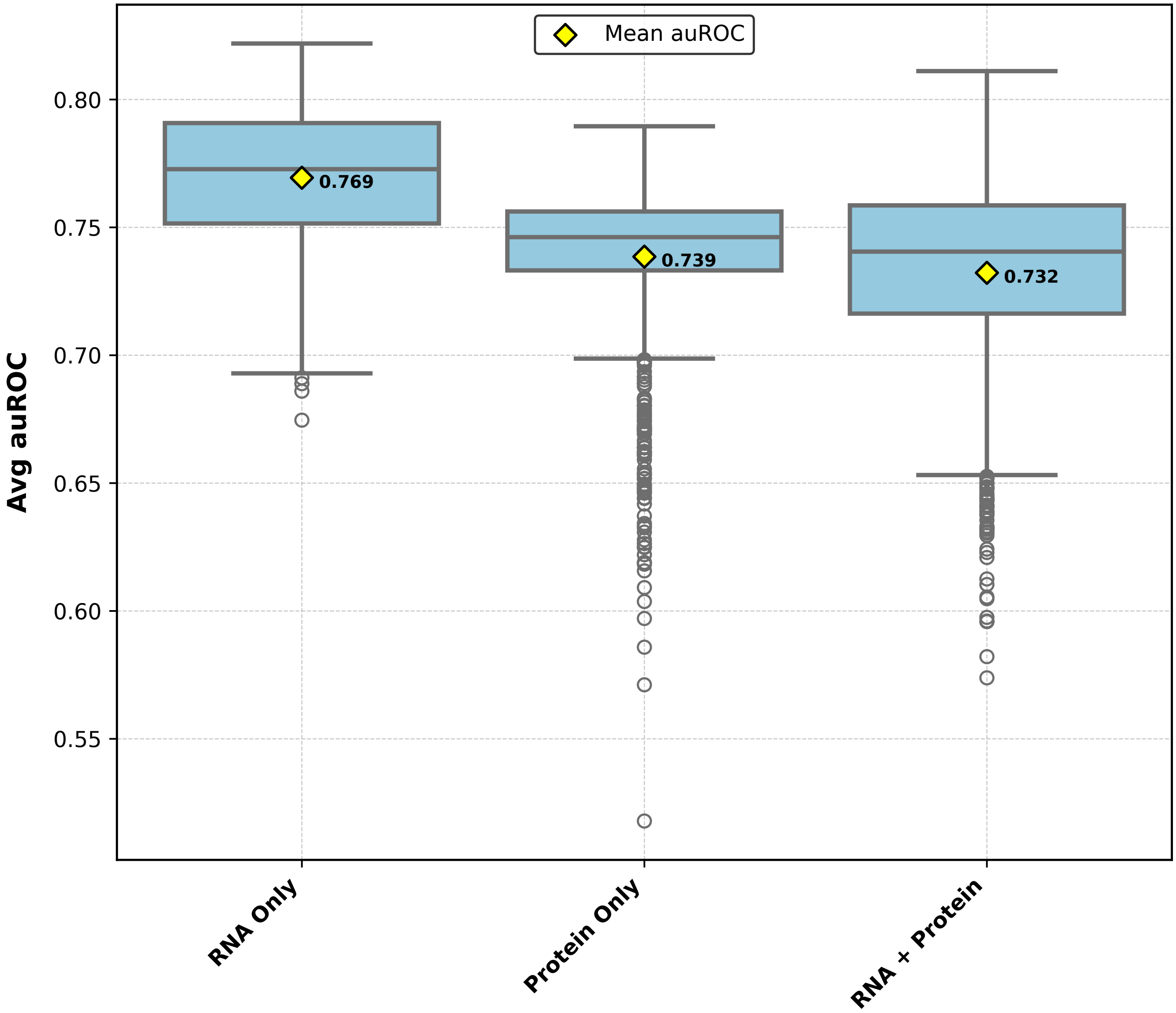

**(B)**

| Model | Avg. auROC | Avg. Accuracy | Avg. Sensitivity | Avg. Specificity |
|---|---|---|---|---|
| mRNA Expression Features Only | 76.9% ± 2.6% | 78.5% ± 5.5% | 76.5% ± 8.6% | 78.7% ± 6.9% |
| Proteomic Profiling Features Only | 73.9% ± 3.1% | 85.4% ± 2.6% | 75.6% ± 3.7% | 86.5% ± 2.9% |
| mRNA Expression + Proteomic Profiling Features | 73.2% ± 3.7% | 80.1% ± 04.7% | 73.2% ± 8.0% | 80.9% ± 6.0% |

**Figure S2:** *(A)* RNA-Seq and proteomic profiling features produce similarly performing models. We formulated three models using a paired 23 tissue RNA-seq and proteomic profiling dataset (Walley et al., 2016) to explore whether RNA-seq or protein abundance features are more predictive of the pollen fitness outcome. The boxplots show mean test-fold auROC from 3-fold cross-validation across 1,000 seeded models; yellow diamonds indicate the overall mean across all seeds. Across seeds, mean auROC was 76.9%, 73.9%, and 73.2% for the RNA-seq only, proeteomic profiling only, and combination models (all 46 expression features), respectively. *(B)* Summary metrics (mean ± SD) for auROC, accuracy, sensitivity, specificity across the same 1,000 seeds.

**Interpretation of Model Coefficients**

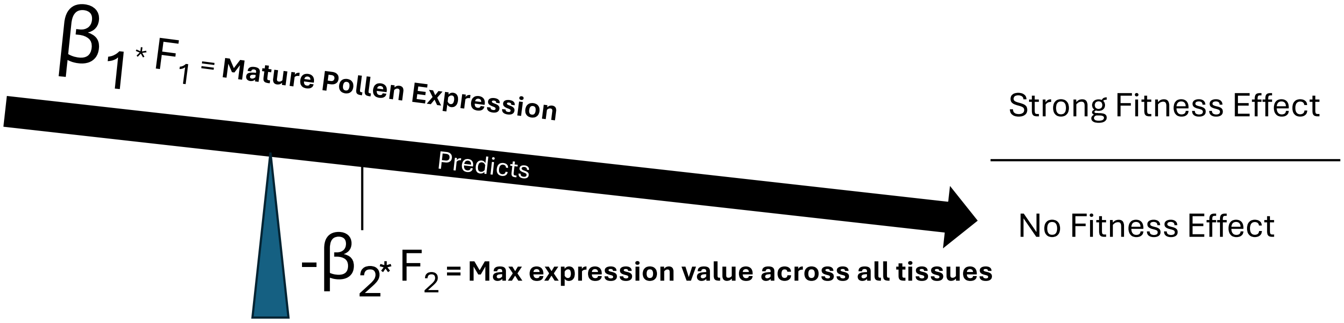

**Figure S3:** Feature importance can be imagined as a weight and balance. In linear models, each feature weight ($\beta$) is a scalar that multiplies the corresponding feature value ($F$) for a given data point. The linear combination of these weighted features determines the predicted probability of belonging to the strong fitness effect class. This process can be visualized as a balance scale, where the magnitude and sign of each weight acts like physical weights tipping a scale. Across many of our early model configurations, the weight ($\beta_1$) of the mature pollen proteomic profiling value and the weight ($\beta_2$) of the maximum proteomic expression values across all profiled tissues (Walley et al., 2016) consistently had similar magnitudes but opposite signs. When mature pollen was the maximally expressed tissue, $F_1$ and $F_2$ were approximately the same value, and because the feature weights were similar in magnitude but opposite in sign, their predictive contributions effectively canceled one another out. Conversely, when a non-pollen tissue had the highest expression, $\beta_2$ dominated, tipping predictions toward a no fitness effect call. This pattern suggested that the gene model expression level in mature pollen, relative to all other maize tissue types, could be predictively important. This motivated us to develop our own quantitative measures of expression specificity across the assessed tissues.

#### 2 Model Feature Specifics

##### Final Models Feature Details

**Table S1:** *(A)* Features included in each of the three models. Reported weights represent the average feature coefficients across models from 1,000 three-fold cross-validation dataset partitions. *(B)* Detailed explanations for each feature.

**(A)**

| Model | Feature | Feature Weight |
|---|---|---|
| Genome Model | Frequency of AA 2–nucleotide sequences (dimer) | 11.50 |
| | Average exon length | 7.85 |
| | Codon usage frequency for GCT | -7.16 |
| | Composition/transition/distribution, distribution: property 1, group 3, residue 75 | -7.42 |
| Expression Model | Log-transformed $PSR_P$ | 6.50 |
| | $NPS_R$ | -11.14 |
| | Maximum RNA-Seq expression | -23.32 |
| Combination Model | Log-transformed $PSR_P$ | 11.41 |
| | Number of variants with high predicted impact | 5.38 |
| | Singleton status from Hufford et al. (2021) | 3.12 |
| | Composition/transition/distribution, distribution: property 1, group 3, residue 75 | -4.00 |
| | Codon usage frequency for GAG | -4.22 |

**(B)**

| Feature | Definition |
|---|---|
| Frequency of AA 2–nucleotide sequences (dimer) | Frequency of AA nucleotide pairs across gene model. Calculated as a raw count with reverse strand complements present (Sen et al., 2023; feng Zhu et al., 2016). |
| Average exon length | Calculated as the sum of all exon lengths across all isoforms of a gene model divided by the total number of exons across all isoforms of that gene model (Sen et al., 2023). |
| Codon usage frequency for GCT | Frequency of alanine codon GCT across gene model. The value is normalized by the total count of all synonymous alanine codons in that gene model (Sen et al., 2023; Xiao, 2014). |
| Composition/transition/distribution (CTD) distribution: property 1, group 3, residue 75 | Composition (C), transition (T), and distribution (D) are three descriptors used to quantify amino acid sequence physicochemical properties: the overall composition of a property (C), the frequency with which the property changes along the chain (T), and the positional distribution of the property across the sequence (D). Seven different properties are considered: hydrophibicity, normalized van der Waals volume, polarity, polarizability, charge, secondary structure, solvent accessibility. For each property, the twenty amino acids are assigned to predefined groups. For example, under the hydrophobicity property, amino acids are classified into the three groups: polar, neutral, and hyrdophobicity. The residue measure is specifically for the distribution descriptor, which defines five positional categories for each property group combination. These categories correspond to the positions (as percentage of sequence length) where the first, 25%, 50%, 75% and 100% of a property group combination are observed. For our models, the CTD feature used (CTDD property 1, group 3, residue 75) represents the percentage of total sequence length required to encompass 75% of all hydrophilic amino acids in the sequence. Thus, a lower value indicates more hydrophilic residues are at the predicted protein amino terminus; a higher value indicates more at the carboxy terminus. (Xiao et al., 2015; Dubchak et al., 1995; Sen et al., 2023). |
| Log-transformed $PSR_P$ | $PSR_P$ (Pollen Specificity Ratio, based on Proteomic data) is a derived feature used to quantify, for a given gene model, the specificity of protein abundance in mature pollen tissue. In our models, this value is log-transformed. Specificity is calculated across protein abundance measurements (Walley et al., 2016). The exact procedure for calculating this feature is described in Methods section 3.3.2 |
| Maximum RNA-Seq expression | This feature represents the maximum RNA-Seq expression value observed across all measured tissues for a given gene model (Sen et al., 2023). |
| $NPS_R$ | $NPS_R$ (Non-Pollen Specificity ratio, based on RNA-seq data) is a derived feature used to quantify the RNA expression specificity of the maximally expressed non-pollen tissue. This value is calculated from RNA-Seq expression values (Walley et al., 2016). The exact procedure for calculating this feature is described in Methods section 3.3.3 |
| Paired status: singleton | A binary feature indicating whether a gene model is a singleton in the maize genome, i.e., no syntelog derived from the ancient whole genome duplication has been maintained at the syntenous position in the genome. Feature values were derived from the work of Hufford et al. (2021), as described in Methods and Supplemental Data S6 |
| Number of variants with high predicted impact | Values derived using the SnpEff tool. SnpEff provides values for assessing the putative impact of a variant (high, moderate, low) on the corresponding protein's function. In our final model, the feature value represents only the count of predicted high impact variants across the maize pangenome for a given gene model (Sen et al., 2023; Cingolani et al., 2012) |
| Codon usage frequency for GAG | Frequency of glutamate codon GAG across gene model. The value is normalized by the total count of all synonymous glutamate codons in that gene model (Sen et al., 2023; Xiao, 2014) |

#### Specificity Feature Visualization

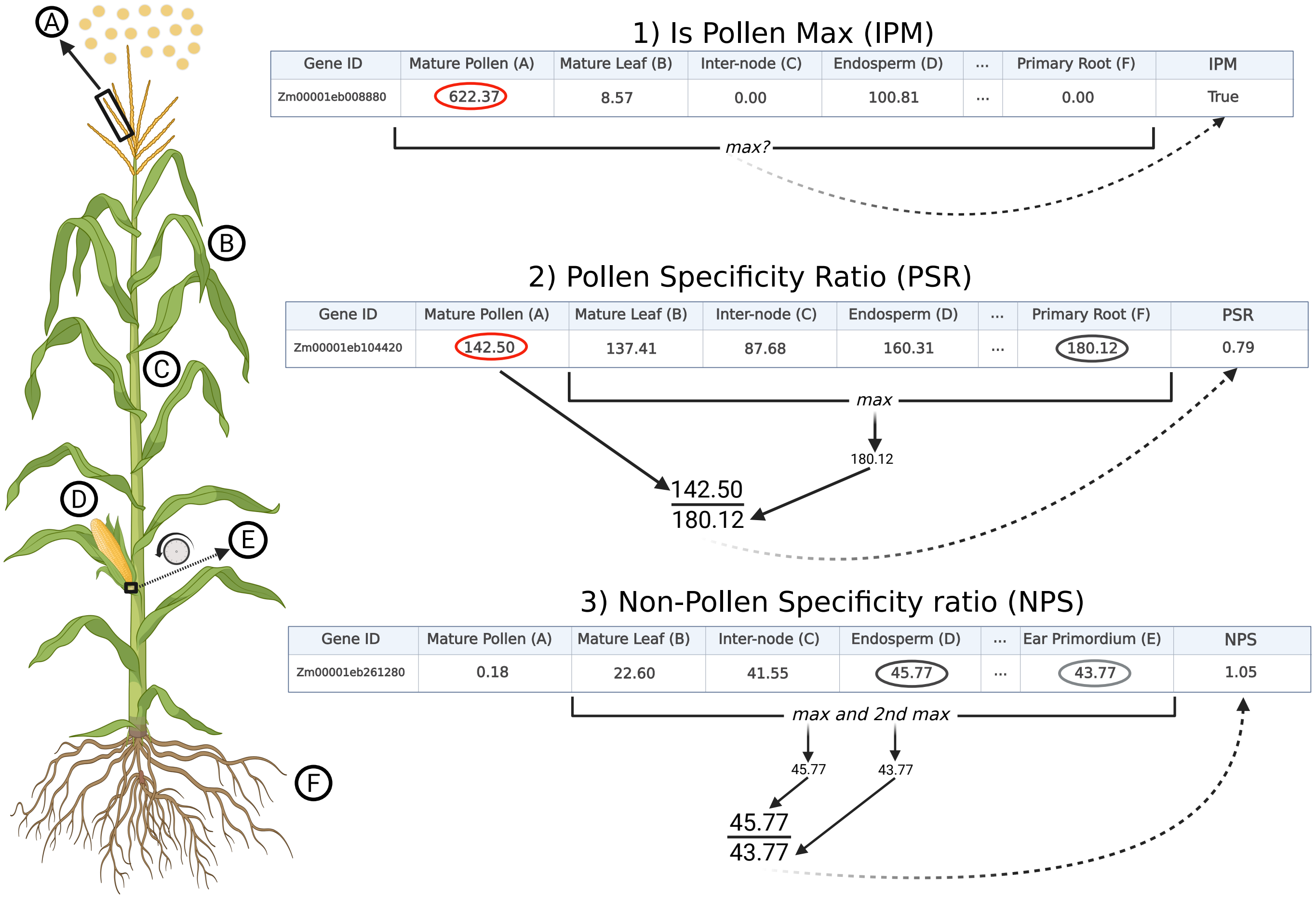

**Figure S4:** Visualization of specificity feature calculations. For each gene model used in training, three tissue specificity features were calculated. Is Pollen Max (IPM) is a binary feature indicating whether the maximum expression value is in pollen tissue. Pollen Specificity Ratio (PSR) is the ratio of pollen expression value to the highest expression value observed among all non-pollen tissues. Non-Pollen Specificity ratio (NPS) is the maximum ratio of expression in any-non pollen tissue to the highest expression value observed among the remaining non-pollen tissues. The calculations shown in the figure illustrate how these features were derived for a single gene model using a subset of tissue types for demonstration purposes. In practice, each of these features was calculated for every gene model in the dataset using expression data (both RNA-seq and proteomic profiling, denoted with an R or P subscript, respectively) from the 23 different tissue types in Walley et al. (2016). Figure created in BioRender. Mueller, S. (2025) `https://BioRender.com/mepxcie`

#### MaizeGDB Features Model Details

**Table S2:** MaizeGDB Features Only Model specifics. Early experimentation models used a collection of RNA-Seq features, proteomic profiling features, and a single genomic sequence feature (gene length).

| Feature | Description |
|---|---|
| `Gene Breadth` | Expression breadth: number of RNA-Seq tissues with measurable expression (Walley et al., 2016). |
| `Anthers R1 Expression Level` | RNA-Seq expression level in anthers (replicate R1) (Walley et al., 2016). |
| `Mature Pollen Expression Level` | RNA-Seq expression level in mature pollen (Walley et al., 2016). |
| `Maximum Protein Abundance` | Maximum protein abundance value (dNSAF) across all tissues (Walley et al., 2016). |
| `Maximum RNA Expression` | Maximum RNA-Seq expression value (FPKM) across all tissues (Walley et al., 2016). |
| `Gene Length` | Gene model length in base pairs (bp). |

#### Expression Feature Subset Test

**(A)**

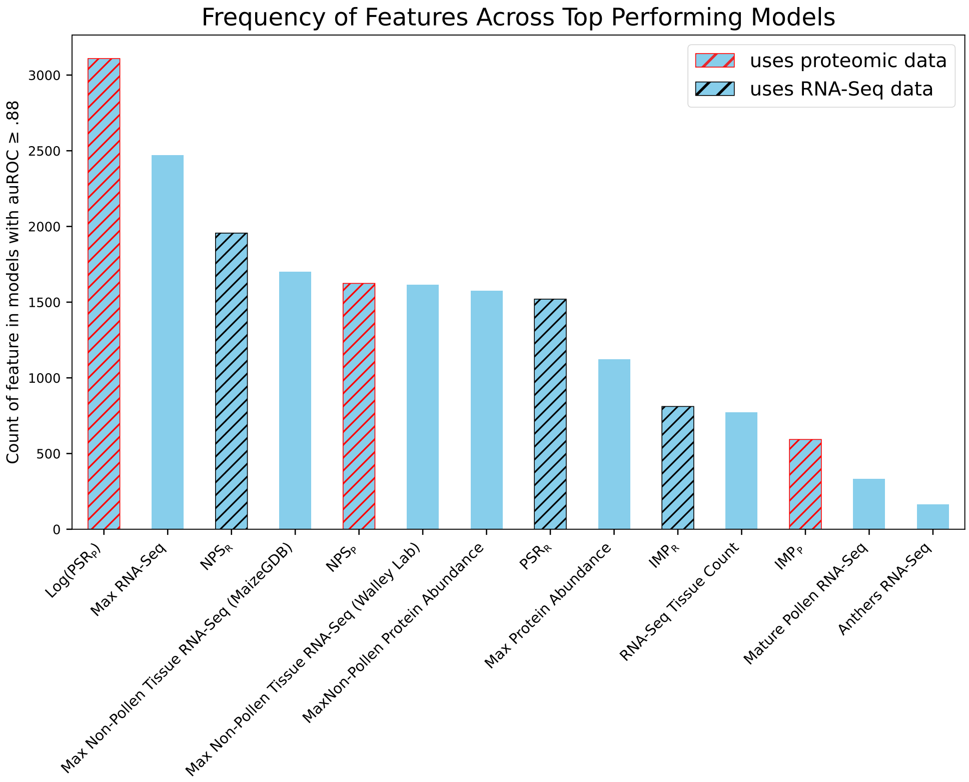

**(B)**

| **Feature** | **Description** |
|---|---|
| `Log Transformed PSR`$_P$ | Log-transformed pollen specificity ratio of protein abundance in pollen to the maximum observed in any non-pollen tissue (dNSAF). |
| `Maximum RNA-Seq` | Maximum RNA-Seq expression value (FPKM) across all tissues (Walley et al., 2016). |
| `NPS`$_R$ | RNA expression specificity within non-pollen tissues: ratio of the highest non-pollen tissue's FPKM to the maximum FPKM of all other non-pollen tissues. |
| `Maximum Non-Pollen RNA Expression` | Maximum RNA-Seq expression value (FPKM) among non-pollen tissues for the large MaizeGDB RNA expression set. |
| `NPS`$_P$ | Protein expression specificity within non-pollen tissues: ratio of the highest non-pollen tissue's dNSAF to the maximum of all other non-pollen tissues. |
| `Maximum Non-Pollen RNA Expression` | Maximum RNA-Seq expression (FPKM) among non-pollen tissues for Walley et al. (2016) dataset. |
| `Maximum Non-Pollen Protein Abundance` | Maximum protein abundance value (dNSAF) among non-pollen tissues (Walley et al., 2016). |
| `PSR`$_R$ | Ratio of RNA expression in pollen (FPKM) to the maximum RNA expression observed in any non-pollen tissue. |
| `Maximum Protein Abundance` | Maximum protein abundance value (dNSAF) across all tissues (Walley et al., 2016). |
| `IMP`$_R$ | True/False that RNA expression is maximal in pollen. |
| `RNA-Seq Tissue Count` | Expression breadth: number of RNA-Seq tissues with measurable expression (Walley et al., 2016). |
| `IMP`$_P$ | True/False that protein abundance is maximal in pollen. |
| `Mature Pollen RNA-Seq` | RNA-Seq expression level in mature pollen (Walley et al., 2016). |
| `Anthers RNA-Seq` | RNA-Seq expression level in anthers (replicate R1) (Walley et al., 2016). |

**Figure S5:** *(A)* Pollen specificity log($PSR_P$) was the most used feature across all top performing expression feature models. A subset search program was run on a collection of fourteen expression features to identify an optimal feature subset. The program tested every combination of these features and ultimately returned the subset used in the final Expression Model. To evaluate whether RNA-Seq or proteomic profiling features were more predictive, both RNA-Seq and proteomic profiling versions of the three specificity measures were included. The bar chart shows the frequency with which individual features appeared among the top performing models (auROC > 88%). This analysis highlighted which features contributed to strong model performance and whether RNA-Seq or protein abundance versions of the specificity features were more informative. *(B)* Detailed descriptions of the fourteen expression and specificity based features.

#### Genome Features Exploration

**Table S3:** Model testing with custom evaluation measurements highlight the predictive potential of genomic features. The table summarizes 15 feature sets sourced from MaizeGDB (Sen et al., 2023) and Hufford et al. (2021), totaling 497 individual features, which were used to explore both genome feature only models and their potential for predictive boost to expression based models. For each feature subset four models were constructed and tested using different combinations of features from the set. Model performance was assessed with three evaluation measurements: *Baseline Predictive Power*† the improvement in feature set performance relative to a 65% auROC threshold; *Downsampling Gain*‡ the difference in performance between full feature set and downsampled feature set; and *Expression Boost*§ the effect on expression model performance after downsampling the feature set. Cell colors denote feature set performance: red = poor, yellow = moderate, green = strong.

| | | | **Genomic Model** | | **Combination Model** |
|---|---|---|---|---|---|
| **Category** | **Feature Subset (# of features)** | **Description** | **Baseline Predictive Power**† | **Downsampling Gain**‡ | **Expression Boost**§ |
| **DNA** | Di-mer (16) | Frequency of 2 pair nucleotide sequences | | | |
| | Tri-mer (64) | Frequency of 3 pair nucleotide sequences | | | |
| | Gene Structure (11) | Gene structural information (e.g. CDS length, Exon number, etc.) | | | |
| | DNA Methylation (9) | CG, CHG, and CHH methylation levels for gene body, upstream, and downstream | | | |
| **Protein** | Codon (136) | Features relating to codons, e.g.: Stacking Energy, GC content, Codon Adaptation Index, etc. | | | |
| | CTDC (21) | Amino acid physicochemical property descriptor that quantifies the overall composition of an amino acid chain with respect to a given property (Dubchak et al., 1995) | | | |
| | CTDT (21) | Amino acid physicochemical property descriptor that quantifies the frequency of transitions between different property-defined groups across amino acid chain (Dubchak et al., 1995) | | | |
| | CTDD (105) | Amino acid physicochemical property descriptor that quantifies the positional distribution of property-defined groups along amino acid chain (Dubchak et al., 1995) | | | |
| | PseAAC (50) | Quantitative representation of composition and sequence order effects of amino acid physicochemical properties (Chou, 2001) | | | |
| | Protein Structure (5) | Six features defining protein structure properties (e.g. Coils, Hotloops) | | | |
| | Subcellular Localization (33) | Protein subcellular localization predictions by WoLF PSORT and DeepLoc | | | |
| **Variation & Evolution** | Insertion Counts (6) | Ac/Ds or UniformMu transposon-based insertion counts calculated for gene body, upstream, and downstream regions | | | |
| | Varionomic (5) | SnpEff variant calls and gene effect prediction | | | |
| | Synteny (8) | Synteny and subgenome mapping, from Hufford et al. (2021) | | | |
| | Miscellaneous (7) | Collection of unique features, e.g., Ka/Ks ratios and gene age | | | |

**Model definitions:**

Model 1 = Full Feature Subset (e.g. all Gene Structure features)
Model 2 = Downsampled Feature Subset (e.g. only highest outcome correlated Protein Structure features)
Model 3 = expression model features + Full Feature Subset (e.g. expression model features + full Codon feature subset)
Model 4 = expression model features + Downsampled Feature Subset (e.g. expression model features + hand-curated highest predictive power Tri-mer features based off of models 1-3).

† **Baseline Predictive Power:** Model 1 or Model 2 auROC gain over baseline 0.65 auROC.

‡ **Downsampling gain:** Downsampling improve auROC between Model 1 and Model 2.

§ **Expression boost:** Model 3 or Model 4 auROC gain over baseline expression model 0.87 auROC.

#### Feature Filter Conditions

**Table S4:** Filter conditions applied to the results of the MaizeGDB and Hufford et al. (2021) feature subset exploration (Table S3). Rows correspond to filter conditions, whereas columns represent the evaluation measures (Table S3) and condition specific notes. For each measure, each cell describes the criteria used for feature selection, based on subset performance (color) and the model type (Models 1-4) from which features were selected. This approach enabled the targeted identification of potential high information yielding features from the larger MaizeGDB and Hufford et al. (2021) feature collections. These composite conditions were ultimately used to define the feature sets incorporated in our Genome Model and Combination Model.

| Condition | Baseline Predictive Power | Downsampling Gain | Expression Boost | Notes |
|---|---|---|---|---|
| 1 | — | — | All weighted Model 4 features for green and yellow subsets | No Ka/Ks features |
| 2 | All weighted Model 2 features for green and yellow subsets | — | — | No dimer features |
| 3 | Two highest magnitude positively weighted and two highest magnitude negatively weighted Model 2 features for green and yellow subsets | — | — | — |
| 4 | All weighted Model 2 features for green and yellow subsets | — | — | — |
| 5 | Two highest magnitude positively weighted and two highest magnitude negatively weighted Model 2 features (included only if weighted across all green and yellow subsets) | — | Two highest magnitude positively weighted and two highest magnitude negatively weighted Model 2 features (included only if weighted across all green and yellow subsets) | — |
| 6 | Two highest magnitude positively weighted and two highest magnitude negatively weighted Model 2 features (included only if weighted across all green and yellow subsets) | — | Two highest magnitude positively weighted and two highest magnitude negatively weighted Model 2 features (included only if weighted across all green and yellow subsets) | Add best performing expression model |
| 7 | — | Two highest magnitude positively weighted and two highest magnitude negatively weighted Model 2 features for green subsets | — | — |
| 8 | — | Two highest magnitude positively weighted and two highest magnitude negatively weighted Model 2 features for green and yellow subsets | — | No Ka/Ks features |
| 9 | All weighted Model 2 features for green subsets | — | — | — |
| 10 | All weighted Model 2 features for green and yellow subsets | — | All weighted Model 2 features for green and yellow subsets | No Ka/Ks features |
| 11 | All weighted Model 4 features for green subsets | All weighted Model 4 features for green subsets | All weighted Model 4 features for green subsets | — |
| 12 | All weighted Model 4 features for green subsets | All weighted Model 4 features for green subsets | All weighted Model 4 features for green subsets | Add best performing expression model |
| 13 | All weighted Model 4 features for green subsets plus the highest magnitude positively weighted and the highest magnitude negatively weighted Model 2 features for yellow subsets | All weighted Model 4 features for green subsets plus the highest magnitude positively weighted and the highest magnitude negatively weighted Model 2 features for yellow subsets | All weighted Model 4 features for green subsets plus the highest magnitude positively weighted and the highest magnitude negatively weighted Model 2 features for yellow subsets | No Ka/Ks; include one high-weighted trimer feature from Model 1 |
| 14 | All weighted Model 4 features for green subsets plus the highest magnitude positively weighted and the highest magnitude negatively weighted Model 2 features for yellow subsets | All weighted Model 4 features for green subsets plus the highest magnitude positively weighted and the highest magnitude negatively weighted Model 2 features for yellow subsets | All weighted Model 4 features for green subsets plus the highest magnitude positively weighted and the highest magnitude negatively weighted Model 2 features for yellow subsets | No Ka/Ks; include one high-weighted trimer feature from Model 1; add best-performing expression model |
| 15 | — | All weighted Model 4 features for green subsets plus the highest magnitude positively weighted and the highest magnitude negatively weighted features for yellow subsets | — | Add best performing expression model |
| 16 | All weighted Model 1 features for green and yellow subsets | The highest magnitude positively weighted and the highest magnitude negatively weighted Model 1 features for green and yellow subsets | — | — |

#### Confidence Analysis of Positive Class Probability Rankings

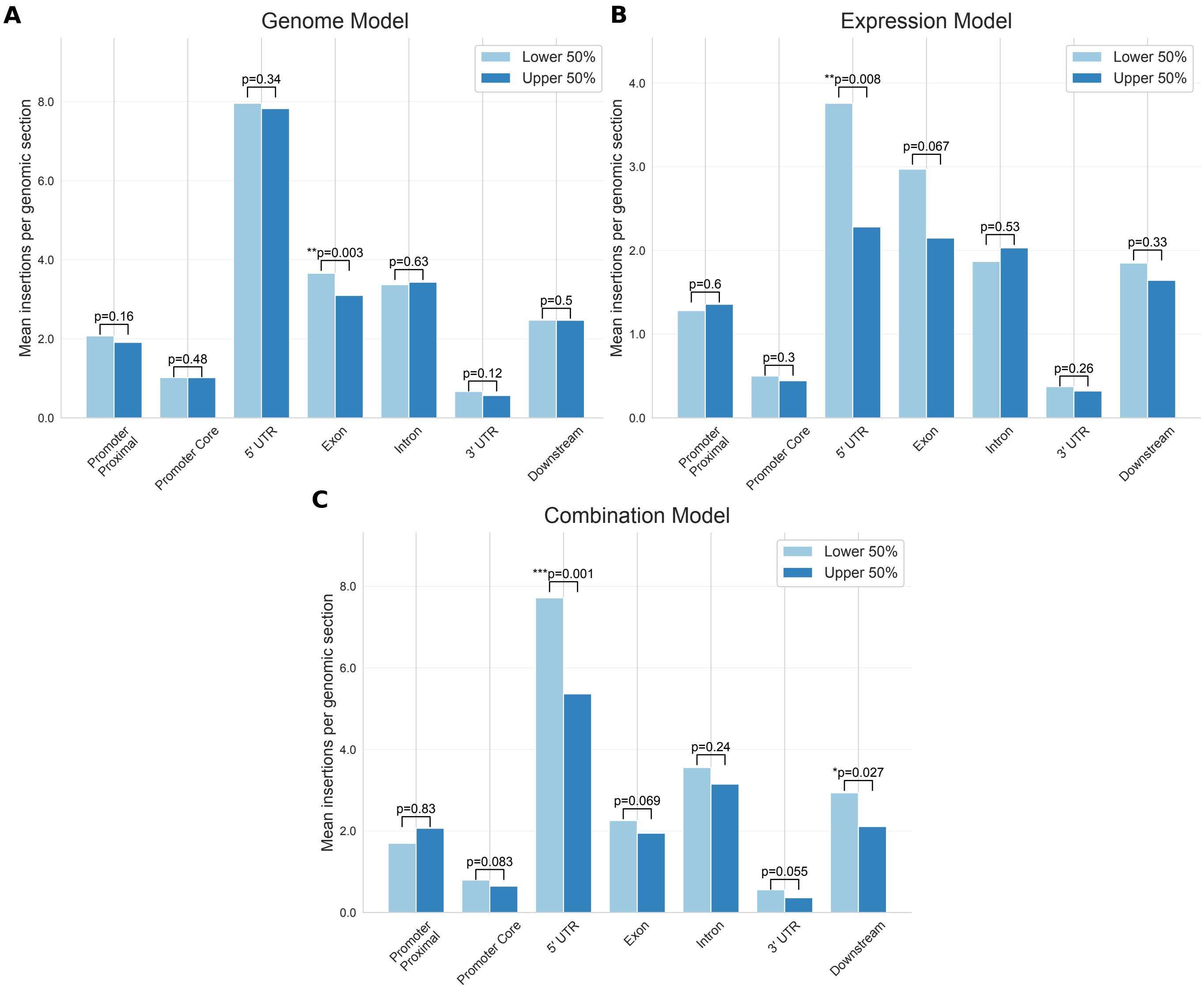

**Figure S6:** Models achieve some success in predicting expected relative frequencies of heritable BonnMu insertions in high probability vs. low probability strong fitness effect gene models. Each panel shows, by genomic section, BonnMu insertion counts for the lower and upper 50% of strong fitness effect gene models (split by model predicted probability) across gene model class predictions. We asked whether model probabilities reflect fitness effect severity – i.e., whether more severe fitness effects receive higher predicted probabilities. If so, we would expect to observe a difference in BonnMu insertion counts between the lower and upper groups of predicted probabilities. To test this, we split predicted strong fitness effect gene models into upper and lower 50% groups by prediction probability. The Genome Model *(A)* showed the expected difference between the two prediction probability groups across all genomic sections except Intron, with a significant difference between the two groups in the Exon section. The Expression Model (*B*) showed the expected difference between the two groups in all genomic sections except for Promoter Proximal and Intron. The Combination Model showed the expected difference in all genomic sections except Promoter Proximal. Notably, permutation testing indicated that both the Expression Model and Combination Model showed a significant difference between the two groups in the Five Prime UTR. The overall number of insertions, and thus statistical power, is highest for this genomic section, due to the known insertion site preference of the *Mutator* transposon near transcription start sites (Zhang et al., 2020).

#### 3 Data and Code

**DataS1_Allele_Specific_Genomic_Information.xlsx**

This dataset contains genomic information for all 305 *Ds-GFP* insertion alleles described in the study. This includes: associated gene model designations, location of the insertion in the gene model (e.g., CDS, intron, 5'UTR, etc), allele-specific primer used for genotyping and Sanger sequencing, and genomic coordinates of the validating flanking sequences.

**DataS2_All_Kernel_Counts.xlsx**

This dataset contains the raw GFP vs non-GFP kernel counts for the 3329 ears used to estimate transmission rates for each *Ds-GFP* allele.

**DataS3_Transmission_Rate_GLM_Final.xlsx**

This dataset contains the GLM-based statistical output for each *Ds-GFP* allele, including estimated transmission rate, adjusted p-value, number of ears assessed for each allele, and the total kernels of each type associated with the counted ears. Based on transmission rate, each allele is assigned to one of three categories described in the main text: strong fitness effect; no fitness effect; and unused (mild or no effect).

**DataS4_Gene_Models_Independent_Alleles_final.xlsx**

This dataset contains information (e.g., transmission rates, location of insertion in gene model, whether the allele information was used for model training) for those alleles representing multiple independent insertions into the same gene models. The alleles are grouped by the fitness categories established in DataS3, and by shared gene model.

**DataS5_Full_Input_Dataset.csv**

This dataset contains the complete set of alleles and associated features used across all model analyses. Subsets of Data S5 were generated for specific model analyses. Data S5 also serves as the primary input file for model training and evaluation in Code S1.

**DataS6_Synteny_Subgenome_Hufford2021.xlsx**

This dataset contains synteny and subgenome information for the entire maize genome, derived from the Hufford et al. (2021) study analyzing the maize NAM founder genome sequences. The initial tab also details the methods used to generate the synteny and subgenome data used for assigning the associated feature values used in modeling.

**DataS7_Model_Training_Datasets**

This file contains the three training datasets used to fit the Genome Model, Expression Model, and Combination Model.

**DataS8_Genome_Wide_Dataset.csv**

This dataset serves as the input for the genome-wide analysis. Specifically, Data S8 includes the larger set of B73 V5 gene models and their associated features. Feature values were sourced from MaizeGDB (Sen et al., 2023) and Hufford et al. (2021), whereas specificity features were calculated using our computational pipeline. This file is required to run the genome-wide analysis in Code S1.

**DataS9_Genome_Wide_Dataset_Predictions.csv**

This dataset contains the same gene models as Data S8, with the addition of predicted class labels and probability values for each gene model generated by the Genome Model, Expression Model, and Combination Model.

**DataS10_BonnMu_Dataset.csv**

This dataset consists of the BonnMu insertion lines and associated measurements used in our analyses, sourced from (Marcon et al., 2020; Win et al., 2024). This file is required to run the BonnMu population insertion frequency analysis in Code S1.

**CodeS1_Project_Code.ipynb**

This Jupyter Notebook contains the analysis code, including the modeling pipeline, genome-wide prediction workflow, BonnMu analysis, cross-validation seed test, and feature subset search analysis.
